# Ultra-high-throughput screening of antimicrobial combination therapies using a two-stage transparent machine learning model

**DOI:** 10.1101/2024.11.25.625231

**Authors:** Margaret M. Reuter, Katherine L. Lev, Jon Albo, Harkirat Singh Arora, Nemo Liu, Shenghao Tan, Madeline R. Shay, Debmalya Sarkar, Aaron Robida, David H. Sherman, Rudy J. Richardson, Nate J. Cira, Sriram Chandrasekaran

## Abstract

Here, we present M2D2, a two-stage machine learning (ML) pipeline that identifies promising antimicrobial drug combinations, which are crucial for combating drug resistance. M2D2 addresses key challenges in drug combination discovery by predicting drug synergies using computationally generated drug-protein interaction data, thereby circumventing the need for expensive omics data. The model improves the accuracy of drug target identification using high-throughput experimental and computational methods via feedback between ML stages. M2D2’s transparent framework provides mechanistic insights into drug interactions and was benchmarked against chemogenomics, transcriptomics, and metabolomics datasets. We experimentally validated M2D2 using high-throughput screening of 946 combinations of Food and Drug Administration (FDA)- approved drugs and antibiotics against *Escherichia coli*. We discovered synergy between a cerebrovascular drug and a widely used penicillin antibiotic and validated predicted mechanisms of action using genome-wide CRISPR inhibition screens. M2D2 offers a transparent ML tool for rapidly designing combination therapies and guides repurposing efforts while providing mechanistic insights.

## INTRODUCTION

Pathogens are becoming progressively drug-resistant, while few new classes of drugs have been discovered^1–3^. In the United States, the Centers for Disease Control and Prevention has documented 2.8 million yearly cases of antibiotic-resistant infections, resulting in more than 35,000 deaths and immense strain on our healthcare systems^1^. Combination drug therapy is increasingly being used to combat drug resistance in cancers and infections^3,4^. Drug combinations have been shown to suppress antibiotic resistance by simultaneously attacking multiple targets^3^. However it is difficult and expensive to engineer a single compound that attacks multiple targets^3^. Even in nature, compounds are produced in combinations by bacteria and fungi to create synergistic multitarget inhibition and microbes rarely produce single compounds^3,5^. Multi-drug therapeutic regimens are currently chosen based on trial and error, which has resulted in some FDA approved combinations that are used clinically such as the synergistic combination of trimethoprim – sulfamethoxazole. However improper selection of drug combination therapy can spread resistance from ineffective exposure^3^. Due to the vast sample size of available drugs, favorable combinations must be predicted using computational models.

Currently, most computational models rely on experimental datasets to predict synergistic combinations, which limits the scale of drugs that can modeled^6–8^. More recently, a deep learning predictive model that used drug – target interaction data as input has been developed to find synergistic drugs combinations for COVID-19 treatment^9^. The drug–target predictions used in the model were mined from experimental databases, limiting the model’s range. Computational models are also most often developed using black-box artificial intelligence methods, where the inner workings of a model are invisible making it uninterpretable. Consequently, they revealed few insights about the biochemical principles behind drug synergy and drug modes of action.

To address these challenges, we have created M2D2 (Mechanistic Machine learning for Drug – Drug interactions), which uses drug-target interactions to predict drug-drug synergy. M2D2 uses a two-stage ML pipeline; the first ML model predicts drug-protein interactions, which are used as inputs to a second ML model that predicts drug-drug interactions. Notably, the first ML stage can utilize any method that infers drug associations. Here we employ multiomics, molecular docking, and a custom ML model as proof of concept. In addition, feedback of feature importance from the second ML model provides further improvement in the accuracy of drug-protein interaction predictions by the first ML layer. The feature importance feedback provides a framework to address a key challenge in drug discovery; currently, identifying drug targets using both experimental omics and computational methods can result in multiple false positives.

The M2D2 model has highly flexible input requirements and can use various experimental omics datasets or computationally generated drug-protein interactions to predict synergy between drugs. By using computationally derived drug–target inputs, the M2D2 model can within an hour predict all necessary drug – target input features and subsequently synergy between drugs. Our model with computationally generated drug–target features matches the performance of models that require experimental data as inputs^6,10^. M2D2 serves as a prototype for highly adaptable, inexpensive, and fast predictive models that use exclusively computational drug-protein features as inputs derived from protein amino acid sequences and drug structures. It has also been structured to be a white-box ML model. Its input features and important features filter can be used to investigate both single–drug and drug-drug synergy mechanisms of action.

As a validation study, M2D2 was used to rapidly generate interactions between more than 2000 FDA-approved drugs to identify repurposed drug combinations against *E. coli. E. coli* was chosen as a model system because it allowed for high-throughput genetic and drug combination screening and has extensive available multiomics and drug interaction data. M2D2 predicted novel synergy for the drug fasudil, typically used to treat brain aneurysms and stroke patients, with mecillinam, a widely used penicillin antibiotic. Importantly, the transparent ML framework behind M2D2 allows users to gain better insight into the biochemical interactions behind synergy. We benchmarked mechanisms of action from M2D2 with chemogenomics, transcriptomics, and metabolomics, and using newly generated CRISPR interference (CRISPRi) screening datasets.

## RESULTS

### Construction of the M2D2 model using multiomics and drug-target interactions

M2D2 predicts synergy between drug pairs or triplets using drug – target interactions as input features in a Random Forest TreeBagger ML model (Figure 2A). To combine two or more drug – target interaction profiles into a single drug – drug profile for prediction, Boolean operations were used to find the union and intersection of drug-target profiles, as done in similar chemogenomics-based models^10^. To allow for flexibility in model construction and comparison between data types, M2D2 was built to use drug – target inputs from multiple omics datasets and computational binding affinity calculations (Figure 2B). The targets in this study include genes, metabolites, and proteins, which correspond to each input dataset (chemogenomics, transcriptomics or proteomics, metabolomics, molecular docking, and ML). The drug-drug interaction training data was taken from various large-scale experimental studies in *E. coli* (Figure 2C). The experimental drug – gene/protein/metabolite interaction features were extracted from drug response chemogenomic^11^, metabolomic^12^, and transcriptomic^13^ data from literature. For all three experimental datasets the highest percentile of both positive and negative z-scores or log10 fold changes were considered when determining if a drug and gene/metabolite had a strong association. A range of cutoffs for inferring association was used to robustly assess ML performance. Molecular docking was used to derive binding affinity and cluster member number, which were both used to determine a strong drug – protein interaction. Each of the five datasets were uniformly processed to infer a level of interaction between drug and protein/gene/metabolite based on each data type.

**Figure 1.**
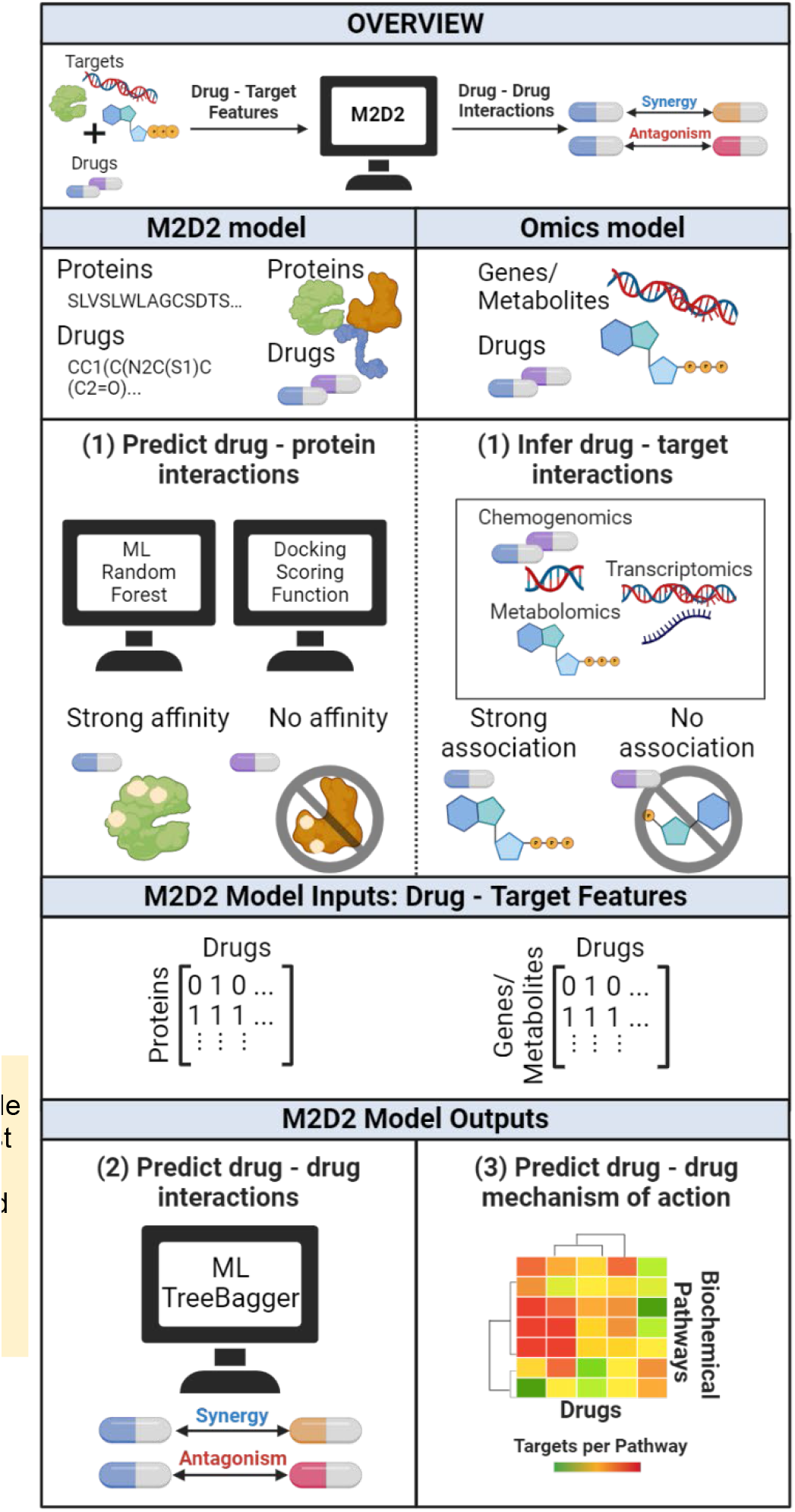
Overall schematic of the M2D2 model design and capabilities. Input features are drug – target interactions. Targets include genes, metabolites, or proteins. The model uses a bagged random forest algorithm to predict synergy, antagonism, or additivity between two or three drug combinations. M2D2 can use either computationally predicted or omics derived inputs features. The datasets the model used as input include target features calculated using molecular docking and machine learning, and features derived from published chemogenomic^11^, metabolomic^12^, and transcriptomic^13^ datasets.

**Figure 2.**
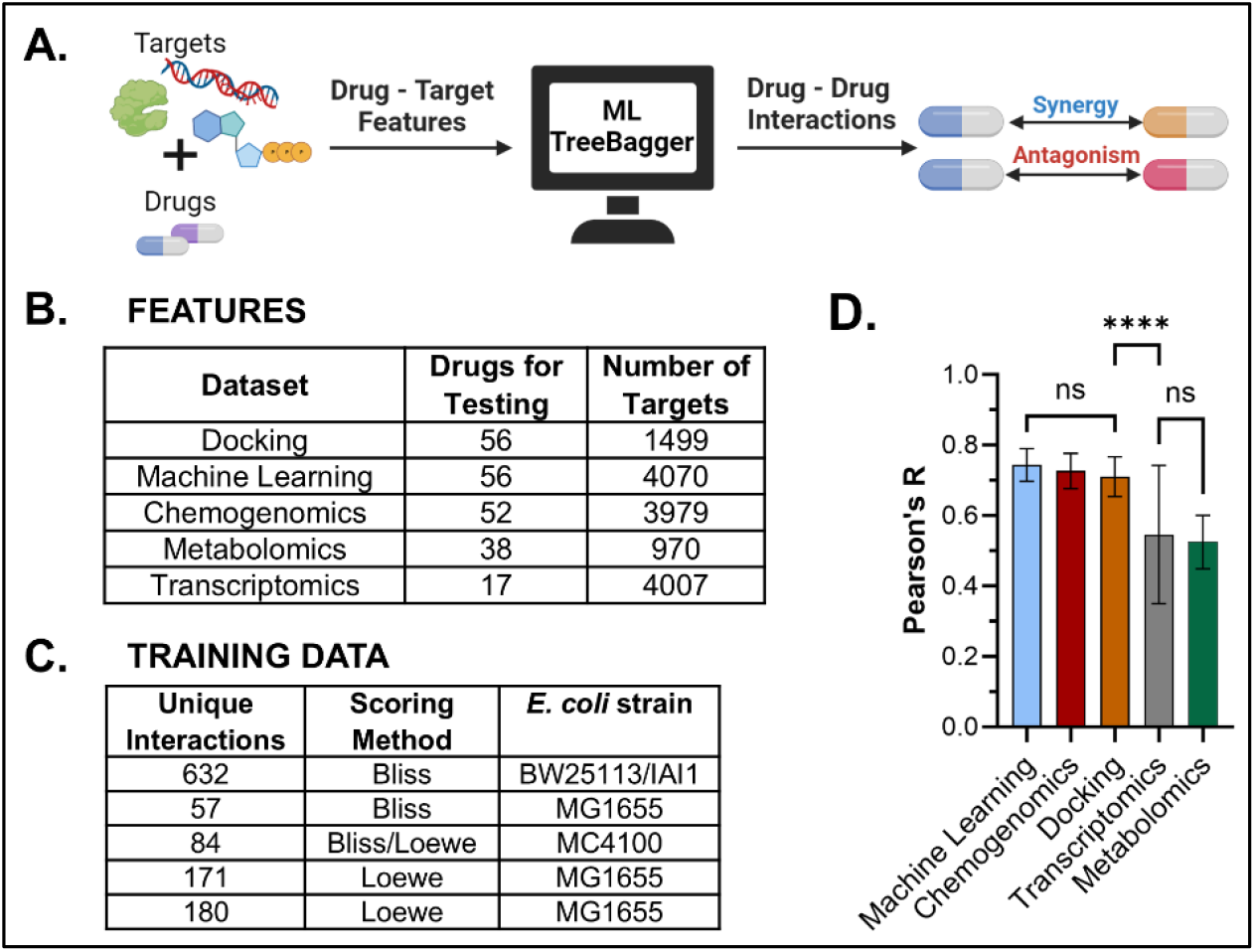
Overall design and performance of M2D2 model in predicting synergistic drug combinations. (A) Simplified schemat the M2D2 model. (B). The number of drugs that were in the dataset a were also in model training data are listed under the Drugs for Testin column. The number of targets includes proteins, gene, or metabolite ry depending on the dataset. (C) Training data for M2D2 was collected 5 different sources^6,10,18–20^. Interactions include primarily two-way combinations and a few three-way combinations. The method for calculating synergy and E. coli strain used in each study is noted. (D) Pearson’s r correlation calculated using 50 iterations of 70/30 hold ou method. The graph is sorted from highest to lowest average Pearson Significant differences between datasets of their aggregated Pearson are noted above the bars using brackets. Three datasets (machine learning, chemogenomics, and molecular docking) have significantly er correlations compared to the other two datasets (transcriptomics and metabolomics). Mean and standard deviation specifics are as follows 0.744±0.046, 0.727±0.050, 0.711±0.056, 0.547±0.196, and 0.525±0. for machine learning, chemogenomics, molecular docking, transcript s, and metabolomics, respectively. Specific p-values are noted in Supplementary Fig. 2.

The first ML portion of M2D2, which predicts interactions between drugs and *E. coli* proteins, was based on the previously developed DeepPurpose algorithm^14^. The basic inputs into the model were drug SMILES structures and protein amino acid sequences. We selected Pseudo amino acid composition (PseudoAAC) for representing the set of proteins, and MACCS encoding were used to represent drugs. MACCS creates 166-bit 2D structure fingerprints that have previously been used to measure molecular similarity between compounds^15^. PseudoAAC has been used in bioinformatics to create discrete numerical protein representations while preserving amino acid sequence information^16^. These encodings were then used in a random forest algorithm to predict drug – protein affinities. The model was trained on interactions from the BindingDB database^17^, which contains 16,702 drugs, 1,659 proteins, and 81,410 proteins. The top encodings for drugs, proteins and the ML architecture were determined based on cross validation accuracy and mean squared error in predicting known dissociation constants Kd (Supplementary Fig. 1).

### M2D2 accurately predicts drug–drug interactions using both experimental and computational drug-target interactions

To assess M2D2’s ability to predict drug-drug interactions using diverse types of input data, we evaluated its performance in a *hold-out* analysis, wherein 30% of the available training data was unseen by the model and used as a test set. The 70/30 splitting of the data was done 50 times to minimize bias to specific drug combinations. The model was evaluated using the correlation between the predicted score of the unseen interactions and their known interaction scores. The 50 correlation scores were then averaged together (Figure 2D).

Training data consisted of 669 unique pairwise and three-way drug combinations among 56 drugs, compiled from five large-scale experimental drug interaction studies^6,10,18–20^. Despite these variable factors M2D2 was still able to predict drug – drug synergy with accuracy higher than previously published models using the same datasets^6,10,21^. Even more promising, the computational drug – protein input features preformed as well as chemogenomic input features (Figure 2D). M2D2 Pearson’s r for 50 holdout iterations was 0.744±0.046, 0.727±0.050, and 0.711±0.056 for ML, chemogenomic, and docking inputs, respectively. Metabolomics and Transcriptomics were limited by data availability to fewer drugs, which may have influenced their overall performance.

A second assessment of M2D2’s ability to predict drug-drug interactions was performed using a *leave-one-drug-out* analysis, wherein all combinations containing a specific drug were left out of the training set and used to test the model. Overall, molecular docking performs the best (Pearson’s r = 0.547, p-value 2.89E-86) across individual drugs, followed closely by both chemogenomics (Pearson’s r = 0.535, p-value 1.55E-79) and ML (Pearson’s r = 0.518, p-value 3.66E-77) (Supplementary Fig. 3). Metabolomics and Transcriptomics had lower correlations but were still highly significant with p-values less than 0.01. Of note, the model performance generalized well even for unseen drug types with distinct modes of action from those in the training set. For example, model performance across all datasets was high for rifampicin and erythromycin, which points to the model’s ability to generalize across drug classes (Supplementary Fig. 3, Supplementary Table 1).

These results suggest that computational input features can rival the effectiveness of expensive experimental input data in predicting drug-drug and drugs-protein interactions. One key benefit of the ML computational dataset is its ability to use any protein with a known amino acid sequence, including essential proteins. Molecular docking could also have the same potential benefit. However, the docking calculations, notably for larger proteins, were too time-intensive compared to ML binding affinity calculations.

### M2D2 can be used to investigate mechanisms of action for individual drugs

We next determined how well the computational drug – target interactions used as inputs in the M2D2 model can predict known drug targets. We focused on 16 drugs that were present in all experimental and computational datasets. These drugs are antibiotics with known mechanisms of action, and these mechanisms can be linked to biochemical pathways in *E. coli* using the Kegg database^22^. All individual drug – target interactions from any dataset used in M2D2 can also be mapped onto Kegg pathways (Figure 3A). Advantageously, pathways maps offer a high-level perspective of interaction networks, facilitating a systems biology analysis of the drug-protein/metabolite/gene interactions. Further, this allowed us to directly compare the performance of computational drug – target interactions with experimental omics methods to identify drug – target interactions by comparing with gold-standard Kegg^22^ (pathway targets) and DrugBank^23^ (specific protein targets) data.

**Figure 3.**
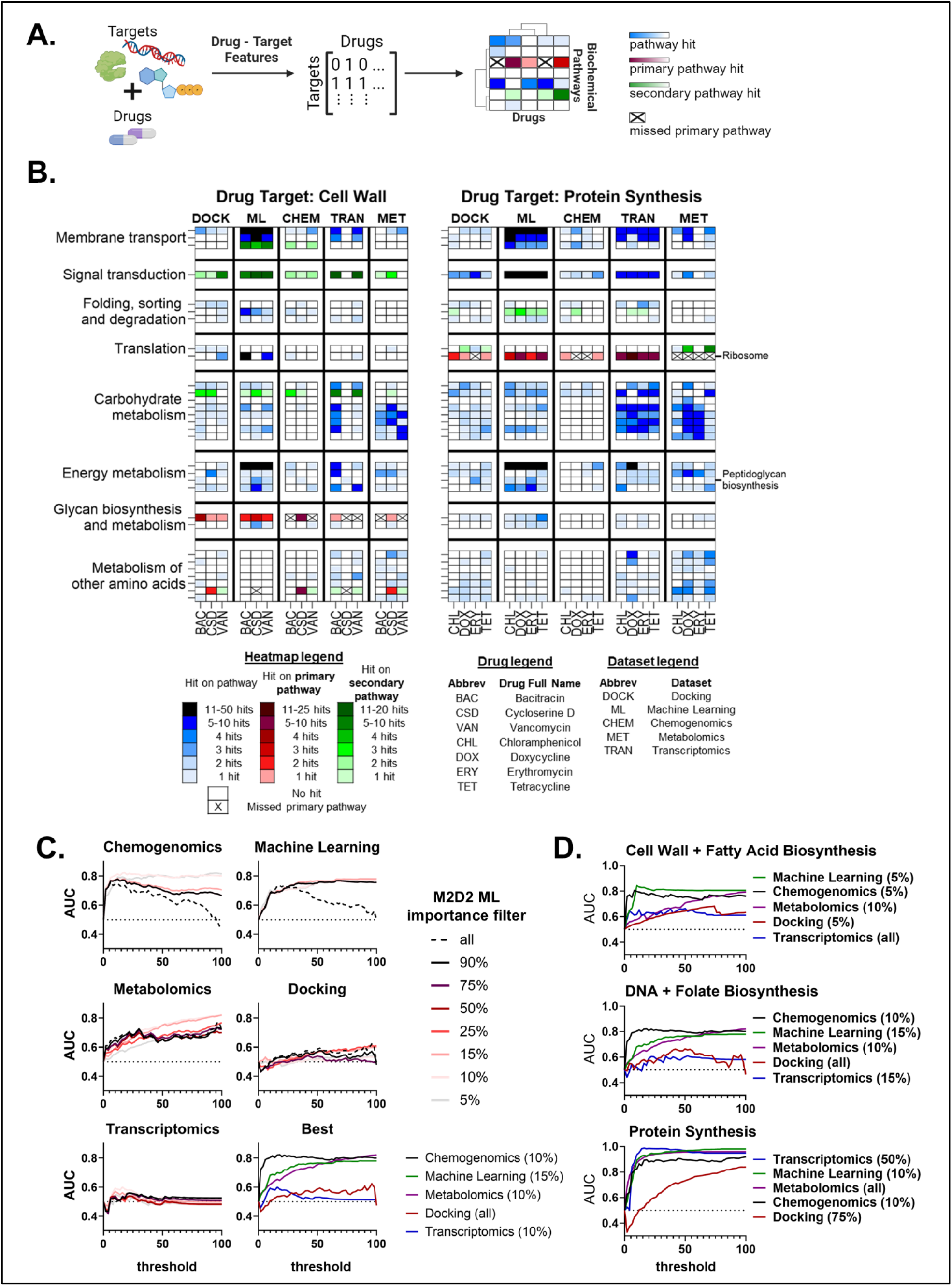
Biochemical investigation of drug mechanisms of action using ML inputs and importance features. (A) Schematic of the overall process. Drug – target features used as input into M2D2 to predict drug – drug interactions were converted into biologically relevant pathways in E. coli. This process was done using annotations from the Kegg database. A strong interaction between drug and target appears as a single hit on a Kegg pathway. Pathways and their respective hits can then be labeled as a primary or secondary pathway for a drug with a known mechanism of action. For example, we would expect erythromycin, a macrolide that inhibits protein synthesis, to hit the Ribosome pathway. (B) Section of the heatmap created to visualize pathways hit by specific drugs grouped by drug class and dataset. Individual pathways we expect a drug to hit for their specific class are labeled on the right of the map. Overarching pathway categories are labeled on the right rather than each specific pathway for readability. This heatmap was created for 50% recall of primary pathway targets. The threshold for a pathway hit was set to reach that recall target to ensure fair comparison. See supplementary GitHub page for corresponding heatmaps for all drugs and pathways. (C) AUC calculation representing a dataset’s ability to hit known drug targets was performed to assess the robustness of the calculation over a range of thresholds used to determine the criterion for a strong interaction between a drug and target. An importance filter was added to the calculation to see if the M2D2 model could improve the AUC calculation. The darker the line the more targets were included in the AUC calculation (e.g., 10% uses only the top 10% most important proteins according to ML). The filter was applied from 0 - 100% in increments of 5%. Not every filter is shown for visual clarity. The best M2D2 ML importance filter was chosen and represented in the 6^th^ graph with the importance percentage in parentheses. (D) The same analysis as panel C was performed but split into specific drug classes. Only the best AUC curves are shown with the corresponding M2D2 ML importance filter percentage in parentheses.

We grouped thousands of drug-protein interactions from various data types into 100 functional pathways. A heatmap of pathways targeted by each drug using information from each datatype was created (Figure 3B). Each pathway in the Kegg database was associated with a list of either genes or metabolites. For the computational datasets, proteins were linked to their respective genes using UniProt^24^ and EcoCyc^25^. If a substantial interaction was found between a drug and target, and that target was associated with a Kegg pathway, that signified one “hit” on a pathway. The area under the precision/recall curve (AUC) was used as a metric to evaluate a dataset’s ability to match known Kegg targets. Overall, chemogenomics, ML, and metabolomics inputs produced the highest AUC values across various “hit” thresholds (Figure 3C). Metabolomics is boosted by the limited number of pathways that could be hit by this dataset, as we excluded pathways that could not be linked to the metabolome. For example, there are no metabolites associated with the ribosome, therefore these pathways were excluded in the recall calculations for the metabolome. The 16 drugs overlapping between datasets were then split into distinct drug classes to evaluate M2D2’s ability to predict mechanisms of action. Nearly all the datasets performed well in the prediction of protein synthesis inhibitors (Figure 3D). In contrast, predicting targets for cell wall and DNA targeting drugs showed lower predictive power across all datasets. ML and chemogenomics features resulted in the best AUC scores across these drug categories.

M2D2 can also be used to investigate specific protein targets rather than higher-level pathways. Interestingly, the transcriptomics dataset appears to be the most robust in an AUC calculation for specific protein targets (Supplementary Fig. 5). ML inputs also performed well, but chemogenomics and docking had much lower AUC values (Supplementary Fig. 5). Metabolomic data were left out of this analysis because specific metabolite targets were not available in Drug Bank, the source for specific protein targets. The decrease in performance of chemogenomics and docking could be attributed to the smaller set of target features available in these datasets (Figure 2B). These datasets might also be lacking essential genes/proteins that are the primary targets of antibiotics. The five datasets were also directly compared with each other using a similar AUC calculation. Instead of comparing them to Kegg, a specific dataset was used to match pathway targets with another data type (Supplementary Fig. 6). M2D2 had the best overlap for chemogenomics, transcriptomics and docking, providing further support for M2D2 predicted targets (Supplementary Fig. 6).

### ML feedback improves drug target predictions

A key challenge in identifying drug targets using both experimental omics and computational methods is the large number of false positives^26,27^. We hypothesized that the second ML model in M2D2 could identify key drug-protein interaction features predictive of drug-drug interactions and thereby reduce false positives. We hence computed feature importance values for all features in each dataset used as input to M2D2. The importance values were taken from the Random Forests algorithm. Feature importance was computed based on the increase in prediction error when a feature was permutated, while the remaining features were unaltered. These values were used to select specific proteins/genes/metabolites of importance, and their corresponding pathways. To quantify the effect of this ML feedback to filter false positive targets we recalculated the AUC values. Promisingly, the importance filter improved both the robustness and overall value of the AUC for all the datasets except molecular docking (Figure 3C). Similarly, at the protein level, the ML importance filter improved performance across datasets, although the overall AUC values were less robust across hit thresholds (Supplementary Fig. 5). At the drug class level, the ML importance filter again improved the performance of nearly all the datasets in almost all cases, except molecular docking for DNA targeting drugs, transcriptomics for cell wall targeting drugs, and metabolomics for protein synthesis targeting drugs (Figure 3D). This indicates that M2D2 is preferentially using drug–target interactions that involve biologically relevant genes, proteins, or metabolites to predict drug–drug synergy. By connecting M2D2 inputs with ML model feature importance, we have created a more accurate and interpretable model.

### High-throughput experimental validation of drug combinations involving repurposed drugs predicted by M2D2

Repurposed drugs present a promising opportunity to rapidly develop new treatments for growing antibiotic resistance. The discovery of entirely new drugs can take up to $3 billion and 15 years, which is often untenable from both economic and public health perspectives^28^. Repurposed drugs are frequently already commercially available, have extensive toxicity profiles, and are FDA-approved, enabling their swift clinical deployment. The availability of prior knowledge on drug structures and interactions, which is typically more abundant for well-studied repurposed drugs, further aids computational drug discovery. Consequently, there is a strong incentive to develop computational tools to identify repurposed drugs for emerging diseases.

Once M2D2’s ability to predict drug–drug interactions using computationally inferred drug-target interactions was established, we used the model to predict interactions for repurposed compounds. We began with a set of 2029 FDA-approved drugs that span a range of treatment purposes from the Broad Institute Repurposing Data Portal^29^. As single compounds, these FDA-approved drugs may not have antimicrobial activity, but we hypothesized that they could potentially boost the activity of traditional antibiotics. Oncology-related compounds were omitted from the set to mitigate the potential for highly toxic combinations^30–33^. The remaining FDA-approved drugs were paired with 56 antimicrobial drugs to generate 113,624 interaction predictions (Figure 4A).

**Figure 4.**
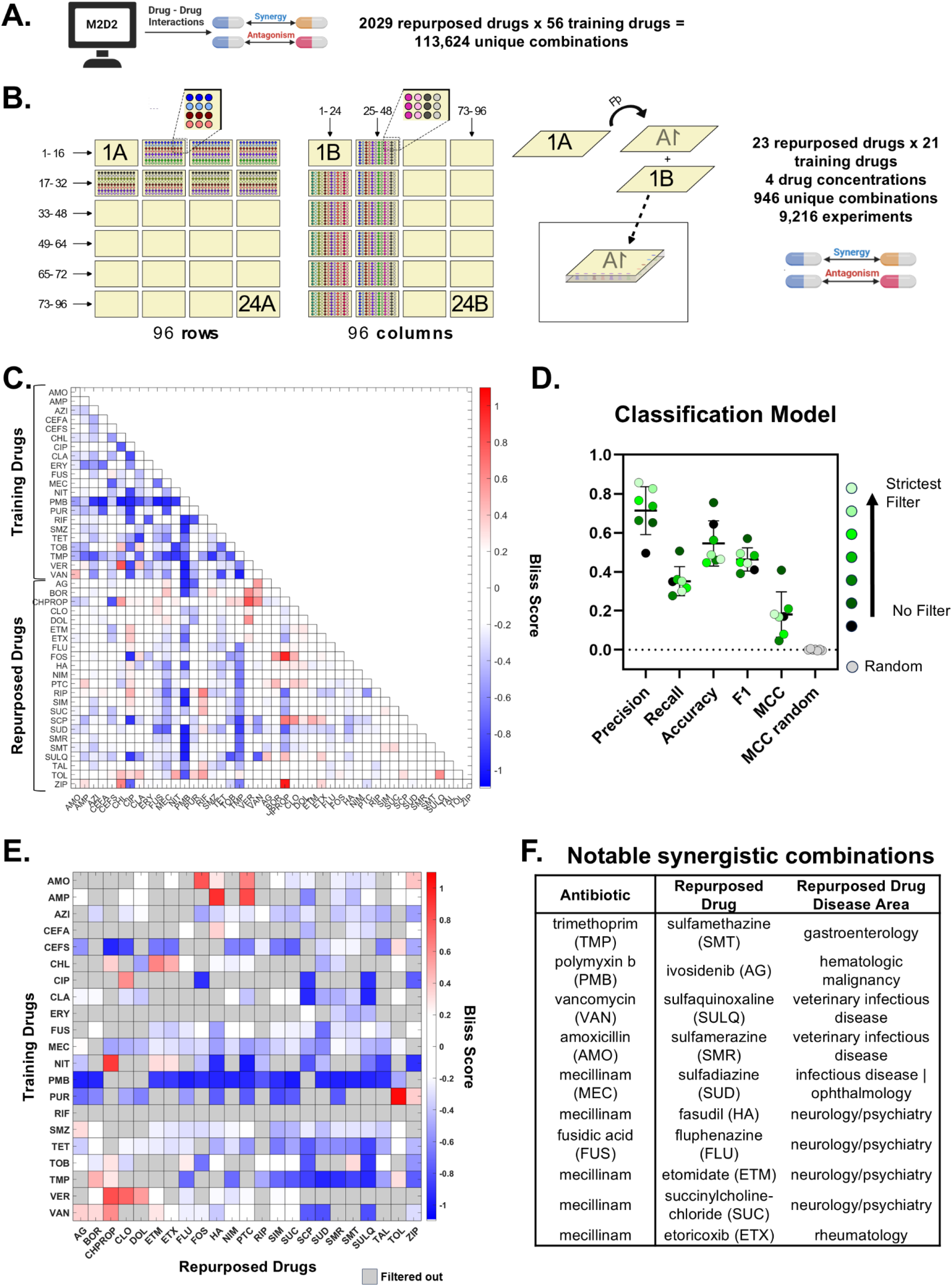
Computational and experimental predictions for interactions of repurposed drugs with traditional antibiotics. The goal was to find repurposed compounds that can boost the antimicrobial power of an antibiotic that is susceptible to antibiotic resistance. (A, B) Overview schematic of both computational and experimental processes, including details of each process. Training drugs refers to the drug compounds used to train the M2D2 model, which were paired with the repurposed drugs to evaluate antimicrobial activity. (B) We performed the high-throughput experiment as an all by all experiment to create four different combinations of drug concentrations and two biological replicates for all experiments. For the full 9,216 combination experiments, we used a SPOTs loader designed to deposit a single liquid in each row or column. The loader can deposit 16 liquids per row or 24 liquids per column. 48 SPOTs plates were used resulting in 24 sandwiches of plates. (C) Results of the experimental high-throughput screening. Only one drug concentration category is pictured for clarity, high dose drug 1 – low dose drug 2. Compounds that are in the set of drug interactions used to train M2D2 are grouped together as are the set of repurposed drugs. The full names of all the drugs can be found in Supplementary Fig. 7. (D) Classification model performance of M2D2 trained on literature training set (Figure 2C). Statistical filters were applied to the experimental data to improve robustness of synergy/antagonism/neutral designation and are noted by color. Mean and standard deviation for each metric are as follows: precision 0.714±0.123, recall 0.352±0.075, accuracy 0.546±0.116, F1 0.463±.060, MCC 0.181±0.116, MCC random 0.000±0.004 (E) Experimental results at the 5^th^ level of filtering also matched M2D2 predictions significantly. The 5^th^ level of filtering corresponds to where standard deviation threshold was met, and classification was consistent across time points and replicates. Filtered out combinations are noted in grey. (F) Table of notable combinations of traditional antibiotic plus repurposed drug resulting in synergy from D. The repurposed drug’s standard usage is noted in repurposed drug disease area.

To confirm some of the computational predictions, 23 repurposed and 21 antimicrobial drugs, were selected for high-throughput experimental screening (Figure 4B). The repurposed drugs spanned a range of intended disease areas (Supplementary Fig. 7), and the antibiotic drugs had a range of mechanisms of action. Repurposed compounds were also selected if they (1) were predicted to show promiscuous synergy, such as sulfachlorpyridazine, which was synergistic with 5 different antibiotics, or (2) had very strong predicted synergies with a single antibiotic such as ripasudil in combination with fusidic acid. Three compounds (bortezomib, nimesulide, and simeprevir) were selected for evaluation because they showed promiscuous antagonism.

High-throughput testing of antibiotics and repurposed drugs was done using the recently developed Surface Patterned Omniphobic Tiles (SPOTs) method^34,35^. Each combination was assessed at two time points and four different drug concentration combinations in duplicate resulting in 9,216 measurements. Bliss scores were then calculated using the growth inhibition data. To provide higher confidence in the SPOTs predictions, statistical filters were applied to the data to obtain consistent synergy, antagonism, or neutral scores over all time points, drug concentrations, and replicates. Of note, this experimental validation dataset is considerably larger than the training dataset used for constructing M2D2.

The predictions from the experimental data were then compared to predictions from the M2D2 model. Notably, M2D2 was trained using the collection of drug interaction scores from literature spanning a range of experimental techniques, *E. coli* strains, and scoring methods. M2D2 was able to match the experimental scores with high precision and Matthews correlation coefficient (MCC) scores, clearly stronger than a randomized set of predictions (Figure 4D). Because the experimental data were collected in a high-throughput method, results were progressively filtered to find the high confidence synergy scores. Stricter filtering resulted in higher precision scores with M2D2, but did not seem to improve recall, accuracy, F1, or MCC (Figure 4D).

M2D2 was then retrained using the experimental dataset to eliminate the variation from multiple literature sources. In this analysis, M2D2 was trained on a subset of the experimental results and assessed on remaining results, but the two sets did not overlap. The retrained model was performed for all four drug concentration combinations, the mean of concentrations, for both time points, and all filtering steps. Overall, the model Pearson’s r scores from a 50 iteration 70/30 hold out were all above 0.58 regardless of filtering and using the mean or median of the concentration scores (Supplementary Table 2). The highest R score was 0.669±0.048 (Supplementary Table 2).

There are numerous predictions from the M2D2 model that were confirmed via high-throughput analysis (Figure 4D). The goal of this screening was to find repurposed drugs that could enhance the efficacy of known antibiotics. Interestingly many of the repurposed drugs that showed synergy in both computational and experimental work were compounds originally approved by the FDA for neurology/psychiatry purposes (Figure 4F). A wide range of non-antibiotic drugs, including compounds that were created for human targets, have been shown to impact a range of bacterial species in the gut microbiome^36^. Antipsychotic drugs, including fluphenazine, showed antimicrobial activity, which was suggested to be part of their mechanism of action involving dopamine or serotonin receptors^36^. Antidepressants have also been shown to provide efflux pump inhibition in both gram negative and positive bacteria^37,38^. All these studies suggest that neurology/psychiatric drugs may have their own antimicrobial effects, although their role in modulating synergy has not been systematically explored previously.

### M2D2 uncovers synergy between fasudil and mecillinam

From the high-throughput screen, we identified a unique combination of mecillinam, a broad-spectrum penicillin, commonly used to treat *E. coli* urinary tract infections, and fasudil, a vasodilator commonly used to treat strokes. Previous work has shown phentolamine, another vasodilator, to be strongly synergistic with macrolides^39^. However, drug interactions between penicillins and fasudil have not been explored. The fasudil/mecillinam combination was then evaluated experimentally using a fractional minimum inhibitory concentration (MIC) assay in a checkerboard. The combination was found to be synergistic for a range of concentrations of both compounds (Figure 5B, Supplementary Fig. 8). Given that fasudil is a repurposed drug, we further evaluated its toxicity. The combination of mecillinam and fasudil was tested for toxicity in both primary liver and kidney cell lines using cell viability assays. It showed little toxicity in either cell line, especially at concentrations that showed synergy against *E. coli* (Figure 5C, Supplementary Fig. 9).

**Figure 5.**
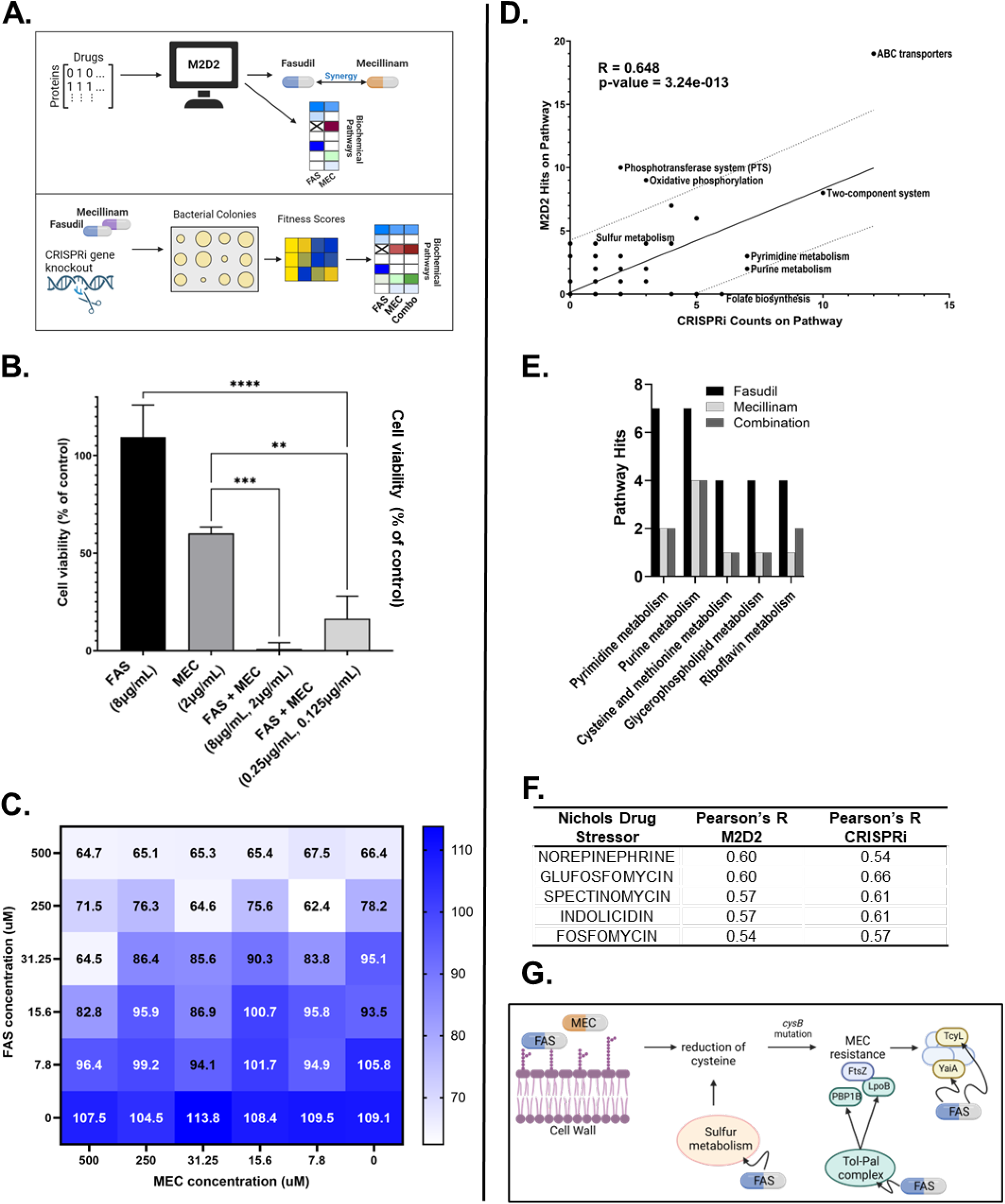
Confirmation of mecillinam – fasudil synergy and subsequent mechanism of synergy analysis using M2D2 and CRISPRi. (A) Schematic of M2D2 and CRISPRi methods for pathway analysis. (B) Fasudil vs. mecillinam Minimum Inhibitory Concentration (MIC) assay in E. coli MG1655 from three biological replicates. P-value significance is noted using an asterisk. P-values were calculated for single drug viability versus combination viability at different concentrations. P-values are from ordinary one-way ANOVA Tukey’s multiple comparisons test in GraphPad Prism 10.4.0. The exact p-values are noted in Supplementary Table 3. The full checkerboard assay for all drug concentration combinations can be found in Supplementary Fig. 8 (C) Toxicity of the fasudil – mecillinam combination at different concentrations in the HEK293 kidney cell line. Lighter shading indicates increased toxicity to the kidney cells. Not all concentrations measured are shown for readability. See Supplementary Fig. 9 for full toxicity profiles in both kidney and liver cell lines. (D) Linear regression of M2D2 pathway predictions with CRISPRi pathway counts. 95% confidence prediction bands are noted in dotted lines and calculated using GraphPad Prism 10.4.0. (E) The bar chart shows pathways hit more by fasudil than either mecillinam or fasudil – mecillinam combination based on CRISPRi data. (F) Top correlated stressors, from Nichols *et al*.^11^ chemogenomic study, with fasudil M2D2 predictions and CRISPRi data from this study. P-values for Pearson’s r values were all less than 1E-8. The top stressors were chosen based on M2D2 correlations. 114 unique stressors, from the Nichols *et al*. dataset, were assessed. More extensive lists of top correlated stressors can be found in Supplementary Table 9 and 10. (G) Potential mechanisms by which fasudil may synergize with mecillinam based on CRISPRi and M2D2 predictions.

We next used the M2D2 model to analyze the mechanism of action for the synergy of the two compounds. Kegg pathways hit by fasudil were compared against other drugs that were synergistic, antagonistic, or neutral with mecillinam in literature and our screening experiments (Supplementary Fig. 10). Hits from M2D2 show sulfur metabolism, pyrimidine metabolism, and ABC transporter pathways as being hit higher in fasudil than the group of drugs antagonistic with mecillinam. These four pathways consistently rank in the top ten pathways at various thresholds for identifying top hits (Supplementary Table 4).

### Chemogenomic investigation of mechanisms of action of fasudil and its synergy with mecillinam using CRISPRi

To validate M2D2’s ability to predict mechanisms of synergy, systematic CRISPR gene inhibition (CRISPRi) was performed for mecillinam, fasudil, and the combination of mecillinam – fasudil. 3802 unique genes were targeted in the CRISPRi assay, which were then mapped to Kegg pathways. Pathway hits were determined by noting genes with greater than 2 or less than -2 log2 fold change in growth after CRISPRi gene knockdown. We performed multiple analyses to parse out mechanisms of synergy, beginning with the analysis of CRISPRi targets alone, comparison of CRISPRi with prior chemogenomic dataset used as input in M2D2^11^ to assess similarity, and performed overlap of CRISPRi targets and M2D2 predictions.

Examining the CRISPRi data on its own, Pyrimidine metabolism, which was prominent in the M2D2 predictions, was shown to have a higher number of hits in fasudil than mecillinam or the combination (Figure 5E, Supplementary Fig. 10). Purine and cysteine/methionine metabolism were targeted more by fasudil than mecillinam. Biofilm formation and phosphotransferase system pathways were consistently enhanced by the combination when compared to fasudil or mecillinam individually (Supplementary Fig. 11).

Similarity among chemogenomic profiles can shed light on potential mechanisms of action of drugs^11,40^. We hence compared the newly generated Fasudil chemogenomics data from CRISPRi with chemogenomics profiles for all the drug stressors from the Nichols et al. study^11^. This study is also the source of chemogenomic data that was used as an input for the M2D2 model (Figure 2B). While fasudil was not a stress condition in the Nichols paper, mecillinam was. We found that the two datasets were significantly concordant. The correlations between Nichols and CRISPRi data for mecillinam were between 0.53 and 0.72 and p-values less than 1E-9. These calculations were performed for CRISPRi log2 fold change cutoff of ±1.5 to 2.5 in increments of 0.5 and Nichols z-score cutoff of ±1.5 or 2.

The Nichols dataset lacks fasudil but contains 114 other unique stresses. The correlation between M2D2 fasudil hits and the 114 conditions in Nichols showed that norepinephrine (Pearson’s r 0.742, p-value 1.08E-18) and glufosfomycin (Pearson’s r 0.603, p-value 3.11E-11) have similar chemogenomic profiles (Figure 5F). Norepinephrine has been shown to be synergistic with antibiotics that require active cell growth for bactericidal effects^41^. Fasudil and norepinephrine also have opposite effects on circulation, fasudil is a vasodilator and norepinephrine is a vasoconstrictor^23,29^.

Similar to fasudil, norepinephrine has also been grouped into the neurology/psychiatry class of drugs, which appears to have multiple predicted synergies with antibiotics by M2D2 (Figure 4D)^41^. While this neuromodulator category can be broad, many subtypes of these drugs have been shown to have antimicrobial activity including anti-neurogenerative disease drugs (fasudil), general anesthesia drugs (etomidate), antipsychotics (fluphenazine)^42^. Generally, there is evidence that drugs preventing neurodegenerative disease, which fasudil has been classified under, impair the biosynthesis of the bacterial cell wall^42^.

Glufosfomycin and fosfomycin also appear in the top five most correlated drug stressors between M2D2, CRISPRi, and Nichols data (Figure 5F). Fosfomycin is an antibiotic that inhibits peptidoglycan biosynthesis through the MurA enzyme^43^. Its mechanism of action involves UDP-GlcNAc and UDP-MurNAc, both of which are fasudil hits in the M2D2 model^43,44^. Fosfomycin has also been shown to enter the *E. coli* cell through two transporter systems, including the hexose-6-phosphate transporter^44^. M2D2 also predicts fasudil hitting the hexose-6-phosphate phosphate antiporter. There is some evidence that fasudil might act on cell wall biosynthesis with similar mechanisms of action to both norepinephrine and fosfomycin. The relationship between these mechanisms and drug combination synergy needs further investigation, however.

### Integrative analysis of M2D2 drug-protein interactions with CRISPRi chemogenomics

When comparing CRISPRi pathway hits to M2D2 predicted pathway hits using both ML models (top 20% of hits from the first ML followed by 15% importance filter from the second ML stage), a strong Pearson’s r correlation was observed (0.648, p-value of 3.24E-13) (Figure 5D). This cutoff and filter were chosen based on the best drug – drug prediction performance and best drug – protein AUC for matching Kegg pathway data across the 16 drugs in Figure 3. Nevertheless, correlations were above 0.5 with p-values less than 1E-8 across cutoffs of 75-90, with and without a M2D2 importance filter (Supplementary Table 5) suggesting that predictions are robust to a wide range of thresholds.

We next linked the CRISPRi data with the ML output to identify high confidence targets. Genes that when knocked down resulted in loss of fitness (i.e., negative log fold change genes from CRISPRi) were compared with binding targets from the M2D2 ML model. This was to investigate whether the M2D2 predicted fasudil hits match any of the genes predicted to increase sensitivity to mecillinam by CRISPRi. Using a cutoff percentile of 80 to determine a strong drug – protein interaction, and a top 15% important feature filter from M2D2, 29 matching genes were found. Genes in sulfur metabolism and ABC transporters were the top pathways in this overlapping gene set (Supplementary Tables 6), suggesting that these pathways might play a role in the synergy between fasudil and mecillinam. Recent literature suggests that a mutation of the *cysB* gene, which degrades cysteine production and metabolism, confers resistance to mecillinam^45,46^. As a sulfur-containing amino acid, cysteine metabolism can be directly linked to sulfur metabolism in *E. coli*^45^. Resistance is attributed to the reduction in cysteine availability and consequently increase in PBP1B, LpoB, and FtsZ proteins. This in turn prompts a 2 to 8-fold increase in 32 unique proteins from oxidative stress, including TcyL and YaiA^46^.

Specific protein hits from M2D2 were examined to provide insights into the mechanism of synergy for the fasudil – mecillinam pair. MreD, RodZ, TcyL, YaiA, CpoB, TolR, and TolA are among the top hits from M2D2 with the ML filter that are also linked to the previously discussed mechanisms of cell well biosynthesis and cysteine metabolism (Supplementary Table 8). MreD and RodZ proteins involved in the rod system, of which PBP2, a known direct target of mecillinam, is also part of^46^. The rod system controls the elongation of rod-shaped bacteria drug peptidoglycan biosynthesis^47^. MreD and RodZ are known to directly interact with MreB during this process^47^. By hitting these proteins, Fasudil could enhance mecillinam’s potency in targeting of the cell wall through the Rod system via parallel pathway inhibition. See the supplementary discussion section for further information on the *cysB* mutation products (TcyL and YaiA), and the Tol-Pal complex proteins (CpoB, TolR, and TolA) which are linked to the PBP1B-LpoB system that is enhanced by the *cysB* mutation.

## DISCUSSION

Here we present M2D2, which predicts drug synergies using computationally generated drug-protein interaction data and circumvents the need for expensive experimentally generated omics data. It is an initial proof of concept for a highly adaptable, inexpensive, and fast predictive model that uses easily available features as inputs. The ML features generated in stage one of M2D2 as features for stage two drug interaction predictions, could feasibly be used with any method that infers drug associations. For this study, we have used multomics, molecular docking, and a custom ML model to both validate M2D2 predictions and demonstrate its flexibility.

We have validated the model using *E. coli* to take advantage of the extensive associated drug interaction and multiomics data, as well as permitting efficient high-throughput genetic and drug combination screening. Conceivably, this M2D2 foundation model could be adapted to more bacteria and perhaps specific bacterial strains given additional drug combination training data using transfer learning techniques^48^. The only other input requirements would be pathogen protein amino acid sequences and drug SMILES structures.

M2D2 has been structured to be a white-box ML model. The use of an ML importance filter elevated the AUC robustness and value across multiple datasets, underscoring the preferential use of biologically relevant interactions by M2D2. An evaluation of predicted drug targets among 100 functional Kegg pathways revealed that ML inputs consistently demonstrated high performance. Overall, these insights highlight the effectiveness of M2D2 in achieving accurate and interpretable drug-target interaction predictions. Its input features and important features filter can be used to investigate both single-drug and drug–drug synergy mechanisms of action.

We demonstrated the utility of M2D2 in narrowing down a huge drug combination sample space to find promising synergy. From the thousands of repurposed drugs combined with a set of antibiotics, over a hundred thousand interaction predictions were made, which would have been unfeasible to comprehensively test via experiments. The synergistic and antagonistic effects of FDA approved drugs on antimicrobials have not been systematically explored previously. Analysis of drugs that were frequently represented among synergistic interactions suggest that neurology/psychiatric drugs may enhance the potency of antibiotics. M2D2 was able to narrow down promising synergies to a testable amount. Several hundred interactions were then validated through a high-throughput screening platform, and the synergistic pair of mecillinam and fasudil was further confirmed via traditional checkerboard assays.

Once the mecillinam – fasudil synergy was confirmed, M2D2 was able to provide biochemical insight into their synergy. Fasudil’s interaction with sulfur metabolism, cysteine/methionine metabolism, and related peptidoglycan biosynthesis pathways suggests multifaceted disruption of the cell wall, enhancing mecillinam’s activity. Fasudil could be targeting proteins in sulfur metabolism that would otherwise lead to mecillinam resistance. M2D2 also predicts fasudil to interact strongly with TolR, another key protein in the Tol-Pal complex, which impacts peptidoglycan biosynthesis. It is possible that Tol-Pal disruption also decreases PBP1B-LpoB functionality, thereby negating the effects of mecillinam resistance created by reduced cysteine production (Supplementary discussion). The Tol-Pal complex has also been linked to pathogenicity in many gram-negative bacteria, making it a promising target for antibiotic development^49^. Overall, specific protein targets predicted by M2D2 seem to indicate that fasudil can facilitate mecillinam’s antimicrobial activity by attacking peptidoglycan biosynthesis both directly and indirectly through the Rod and Tol-Pal systems. Fasudil may also target proteins related to cysteine biosynthesis. This specific set of protein targets could prevent any conferred resistance to mecillinam due to decreased cysteine levels.

One of the main limitations of M2D2 is the need for extensive drug interaction training data. Derived from multiple sources using both Bliss and Loewe synergy scoring methods, the training data has substantial variability and scaling issues. While M2D2 performs better than previously published models, its performance could be improved by using a standardized set of high-quality drug interaction training data, as demonstrated here with the newly generated dataset of 948 drug combinations. While currently implemented for the model organism *E. coli*, as a proof of concept, the M2D2 framework could be extended to different microbial strains in future iterations using orthology-based techniques used in prior chemogenomics-based ML models^10^. A strain-specific model could be readily developed by accounting for differences in protein sequences in the drug – protein interaction calculations. Once a strain model method is validated, it could enable rapid adaptation to new resistant strains as they are being discovered. From a drug discovery perspective, M2D2 could be designed to predict the synergy between antibiotics and novel compounds, such as repurposed drugs or natural products, based on their chemical SMILES structure. The primary limitation to this avenue of advancement would be the protein-ligand modeling for ML features. Most of the BindingDB data used for training involves drug and druglike compounds. If the compounds are too different from the training data from BindingDB, compound–protein binding affinity predictions might be inaccurate. Overall, M2D2 provides insights into drug mechanisms, guides repurposing efforts, and offers a transparent mechanistic ML tool for designing combination therapies.

## METHODS

### ML features: computational datasets

Drug – gene/protein/metabolite interaction scores were calculated using computational methods or extracted from omics literature. *In silico* calculations predicting interactions between 58 drugs and *E. coli* proteins were performed using (1) molecular docking and (2) ML methods. Combined, there are 323,002 completed binding affinity calculations. Molecular docking was performed using VINA and the AMBER14 force field in YASARA version 20.10.4^50,51^. 3-D conformers (2-D conformer if no 3-D conformer was available) of the drug ligands were downloaded from PubChem and energy minimized in YASARA before being used in the docking simulation. Drug ligands were chosen to maximize the overlap between computational and experimental datasets. The set of proteins used for computational modeling were collected from the UniProt database^14^. A protein and corresponding crystallographic structure were selected based on association with *E. coli*, independent of strain, and having resolution less than 2.5Å. The same set of proteins was used in both deep learning and molecular docking methods. Each ligand was docked with each of the 1499 unique *E. coli* proteins 25 times and kept flexible. Docking targets with clustered according toa 5.0Å cutoff. Binding affinity and number of members per cluster were both to determine a strong drug – protein interaction.

Binding affinity was calculated using a model based on the previously developed DeepPurpose algorithm^15^. The basic input into the model was drug SMILES structures and protein amino acid sequences. Encodings for both drug compounds and proteins were tested and selected. For drug encodings Pubchem substructure – based fingerprints, Morgan fingerprints, transformer encoder on ESPF, and MACCS were tested. Transformer encoder on ESPF, Pseudo amino acid composition (PseudoAAC), and conjoint triad features were tested for protein sequences. MACCS were used to represent drugs and PseudoAAC were used for the 4070 set of proteins. MACCS create 166-bit 2D structure fingerprints that have previously been used to measure molecular similarity between compounds^16^. PseudoAAC has been used in bioinformatics to create discrete numerical protein representations while preserving amino acid sequence information^17^. These encodings were then used in a random forest algorithm to predict drug – protein affinities. The model was trained on interactions from the BindingDB database^11^, which contains 16,702 drugs, 1,659 proteins, and 81,410 proteins. This dataset was filtered to remove duplicate interactions and labeling errors, which resulted in 52,142 higher confidence drug – target pairs that were then used to train the ML drug protein interactions model. To validate the drug interaction model, 10-fold cross validation was used (Supplementary Fig. 1).

### ML features: experimental omics datasets

To create a multiomics approach, literature sources were collected for chemogenomic^11^, metabolomic^13^, and transcriptomic^22^ data, while maximizing the number of overlapping drugs between datasets. 16 drugs were present in all three datasets, and they became the primary focus of further analysis. This set of 16 is also in the computational datasets. For all three experimental datasets the highest percentile of both positive and negative z-scores or log10 fold changes were considered when determining if a drug and gene/metabolite had a stong interaction. A false discovery rate cutoff of 0.05 was also used for the transcriptomics dataset. Each of the five datasets were processed to infer a level of interaction between drug and protein/gene/metabolite based on each data type. Z-score or percentile cutoffs were varied and applied to each computational and omics dataset to determine an optimal cutoff for (1) the ML algorithms used to predict drug interactions and (2) a desired recall of known drug targets. These cutoffs were calculated per drug. The result of applying the cutoff was a simplified binary matrix of interaction or no interaction.

### Mechanism of action analysis

Interactions between drugs and the specific gene/metabolites/proteins in each datasets were linked to their *E. coli* pathway maps from the Kegg database^24^. Pathways maps create a high-level view of interaction and reaction networks, which allows for a systems biology analysis of the drug – protein/metabolite/gene interaction matrix. Rather than assessing thousands of small molecule interactions, 100 functional pathways were used. The Kegg database contains 113 pathways for *E. coli*, but 13 pathways were eliminated because they contained no genes or metabolites from any of the omics or computational datasets. Each pathway in the Kegg database was associated with a list of either genes or metabolites. For the computational datasets, proteins were linked to their respective genes using UniProt^25^ and EcoCyc^23^. If a strong interaction was found between a drug and target, and that target was associated with a Kegg pathway, that signified one “hit” on a pathway.

Heatmaps of hits on pathways by each drug were created for each dataset. In addition to simply creating heatmaps of hits per pathway, known drug target pathways were also collected and denoted within the heatmaps. Drug targets were primarily collected from the Kegg database, but if not target was found there, DrugBank^23^ targets were used instead. Targets were then associated with primary and secondary pathways. Primary pathways being directly linked to a drug target, e.g., doxycycline inhibits protein synthesis, the 30S ribosomal unit, so its primary pathway is the Ribosome (eco03010 in Kegg) within the class Translation. Its secondary pathways are Protein Export (eco03060 in Kegg), which is closely associated with the ribosome in the Kegg database and Aminoacyl-tRNA biosynthesis (eco00970), which is associated with the 30S subunit and denoted as a doxycycline target in DrugBank. The final set of heatmaps was created based on 50% recall of the Kegg primary targets for each dataset. Pathways that contain no genes or metabolites on them were ignored, e.g., the Ribosome pathway does not contain any metabolites, thus this pathway was not considered for the metabolomics dataset when calculating recall.

### Evaluation of dataset ability to match known targets (AUC calculation)

To assess each dataset beyond its ability to predict drug – drug synergy, we calculated how well each dataset could infer known drug targets from the Kegg database^50^, via an AUC calculation. An array of known Kegg targets was created for each dataset. Only primary pathways from Kegg were used in the AUC calculations. A cutoff percentile was used to designate hit or no hit for each target. Target hits were then translated into Kegg pathway hits. An AUC was calculated for each percentile using the perfcurve function in Matlab R2021b^52^. The AUC calculation compared how well the binary matrix of hits from a dataset matched the array of known Kegg targets. The same AUC calculation was used to test the dataset abilities to match known specific targets. The Kegg primary pathways were replaced with DrugBank targets^23^. Then the AUC calculation was performed for the binary matrices of each dataset over all percentile cutoffs compared to the specific protein targets.

### ML importance filter

The 2^nd^ ML model in M2D2, which predicts drug – drug interactions, provided feature importance values for each feature in each dataset. The importance values were taken from the Random Forests permutation importance (TreeBagger function in MATLAB R2021b)^52^. These values were used to select specific proteins/genes/metabolites of importance, and thus pathways, for each AUC calculation. The percentage of important elements varied from 5 to 95%. For each percentage of important features, the percentile cutoff that determined a hit varied from 0 to 100.

### ML for drug – drug synergy predictions

Drug – gene/protein/metabolite interaction scores were used as input into the M2D2 model. Single drug – target affinities were binarized using an optimal cutoff percentile or z-score cutoff. The cutoff for a strong interaction/association was varied to get the best ML performance in the training set. Individual Drug – target profiles were combined into drug – drug profile using Boolean operations previously developed in the INDIGO and MAGENTA algorithms^10^. The Boolean operations capture both similarity and uniqueness of the two sets of drug – target profile by calculating their union (sigma score) and intersect (delta score)^6,10,18–20^.

Built in MATLAB R2021b^52^ functions for decision trees, random forest, TreeBagger (a bagged random forest algorithm), and support vector machines were investigated. A regression model was used to predict specific interaction scores. Training data consisted of 669 unique combinations among 56 unique conditions from 5 sources^34,35^. Training and testing sets of drug interactions were split 70/30 randomly for a holdout calculation. For each algorithm, 50 iterations of the holdout calculation were performed, and Pearson’s r correlation was calculated. The interactions that were in common between sources were averaged together. Bliss and Loewe scoring methods and *E. coli* strains were all used in the same model.

For the leave-one-out analysis, a single drug was chosen, and all interactions involving that specific drug were left out of the training set. Pearson’s r was calculated for predictions of the test set which included all the drug interactions with that specific drug. Only drugs involved in at least 25 drug – drug interactions were used to ensure an adequate amount of training and testing data available.

### High-throughput experiments

High-throughput testing of antibiotics and repurposed drugs was performed using the Surface Patterned Omniphobic Tiles (SPOTs) platform^34,35^. We first plated each drug to determine the MIC. First, a total of 44 drugs (23 repurposed and 21 training drugs) were loaded onto SPOTs plates using different sized spots to have different volumes of each drug. On a separate SPOTs plate, *E. coli* diluted to 2^*^10^6 CFU/mL was loaded at constant volumes. These two plates were sandwiched together, and inhibition was determined by using dark-field microscopy to measure an increase in light intensity, indicating cell growth. These images were taken at 8, 12, 16 and 24 hours and were used to create growth curves. We then fit a hill equation and used this to extract MIC20 (high) and MIC80 (low) for each drug’s high and low concentrations.

For the full 9,216 combination experiments, we used a SPOTs loader designed to deposit a single liquid in each row or column. Each drug was used at two drug concentrations which resulted in 88 loaded drug solutions. We also included four positive controls (containing *E. coli* and LB broth) and four negative controls (containing LB broth). We performed this experiment as an all by all experiment to create four different combinations of drug concentrations and two biological replicates for all experiments. The four different combinations of drug concentrations consisted of: high concentration of drug 1 combined with high concentration of drug 2, high concentration of drug 1 combined with low concentration of drug 2, low concentration of drug 1 combined with high concentration of drug 2, low concentration of drug 1 combined with low concentration of drug 2. High and low drug concentrations being MIC20 and MIC80, respectively. We mixed *E. coli* with each drug dilution right before plating to achieve a final concentration of 1^*^10^6 CFU/mL. All liquid handling steps were performed in a cold room to reduce any interactions and decrease any evaporation. We used a SPOTs loader that can load 16 liquids per row or 24 liquids per column, and slid from plate to plate to decrease loading time. We used a total of 48 SPOTs plates which resulted in 24 sandwiches of SPOTs plates and 9,216 experiments. These plate sandwiches were sealed with paraffin wax around the edges and incubated at 37°C. Dark field measurements were taken at 8 and 16 hours. Raw Bliss scores were scaled based on maximum synergy or antagonism values calculated using the fraction of dead cells.

To provide higher confidence in the SPOTs predictions, statistical features were applied to the data and was organized in the following structure: (1) time point one, (2) time point 2, (3) standard deviation of each drug – drug combination less than 5000 for time point one, (4) standard deviation of each drug – drug combination less than 5000 for time point two, (5) consistent synergy, antagonism, or neutral scores over both time points, (6) same criteria as 5 and a standard deviation in the single drug MIC was less than 5000, (7) same criteria as 6 and multiple concentrations show the same interaction category (synergy/antagonism/neutral).

### Traditional checkerboard assay

#### Bacterial Strains, Compounds, and Reagents

The *E. coli* MG1655 strain used in this work was provided by the USDA-ARS Culture Collection (NRRL). Fasudil and mecillinam powders were purchased through Cayman Chemical (Ann Arbor, USA) and stored in powdered form at -20oC prior to resuspension in sterile ddH2O for experimental use.

#### Checkerboard Assay Protocol

Fasudil and mecillinam synergy was determined in *E. coli* MG1655 using a checkerboard assay. Briefly, each compound was serially diluted 2-fold in 50μL in two 96-well flat-bottom microtiter plates at a starting concentration at 2 x MIC for each compound. The first compound was transferred to the corresponding wells containing the second compound such that every row contained serial dilutions of a compound, and every column contained serial dilutions of the second compound, in the perpendicular direction. Once completed, 50μL of LB medium was added to every well, following which 50μL of an overnight culture diluted to 0.08-0.1 OD600 was added to every well besides the media blanks. Each plate additionally contained each single compound without the addition of the adjuvant compound diluted 2-fold. Plates were incubated at 37°C for 18-24 hours and read at OD600. Experiments were completed in triplicate.

### CRISPRi experiments

#### Bacterial Strain and Plasmid Library

*E. coli* genome-wide inhibition library was a gift from David Bikard (Addgene #115927). LC-E75 was a gift from David Bikard (Addgene plasmid # 115925). Strains were cultured on Luria-Bertani Broth (LB, BD Biosciences, USA) supplemented with 12g agar for solid plates.

#### Preparation of Electrocompetent Cells and Library Transformation

Briefly, an overnight culture of LC-E75 was diluted into 1 L of LB medium and incubated in an orbital shaker at 37°C until reaching an OD600 of 1. Cells were pelleted at 4°C and washed three times with chilled 10% glycerol in ddH2O, centrifuging in between washes at 1000g for 15 minutes. Electrocompetent cells were aliquoted into pre-chilled Eppendorf tubes and frozen at -80°C. To prepare cells for library transformation, thawed electrocompetent cell aliquots were suspended with 50 ng of the purchased plasmid library (see above) in a pre-chilled 1 mm electroporation cuvette and electroporated with the following settings: 1.8kV, 25uF, 200Ω. Following electroporation, 980μL of Super Optimal broth with Catabolite repression (SOC) medium was added to the cuvette and pipetted to resuspend prior to incubation for 1 hour with shaking at 37°C. The transformation mixture was serially diluted 6-fold 1:10 and spotted on an LB plate with kanamycin added to quantify the transformation. The remainder of the cells were diluted in 10mL of SOC recovery medium and plated on 145mm round LB-kanamycin agar plates. All plates were incubated overnight at 37°C, and transformation efficiency was quantified the following day. Aliquots of transformed cells were stored in glycerol stocks at -80°C.

#### Chemogenomic CRISPRi Screen

Transformed, electrocompetent LC-E75 cells were thawed and 5μL per sample were diluted in 100mL LB with kanamycin to achieve a starting OD600 of 0.001. Cells were grown for approximately three hours to achieve an OD600 of 0.2 and 50mL of culture was sampled for the time 0 sample prior to dCas9 expression. Following, all flasks were treated with 25μL of a 1 mg/mL stock of anhydrotetracycline (aTc, Cayman Chemical, USA) to induce dCas9 expression. Cells were grown at 37°C until an OD600 of 2, at which time 5mL of culture was sampled for the post-induction timepoint. Cultures were diluted into 100mL of LB with aTc and kanamycin added and allowed to grow to an OD600 of 0.4, at which point 20mL of culture was collected for a t0 timepoint. Following, cultures were treated with sub-inhibitory concentrations of fasudil, mecillinam, and fasudil + mecillinam and grown for 2 hours at 37°C. Cultures were pelleted and library was immediately extracted by miniprep (Qiagen, USA). All samples and controls were collected in triplicate.

#### Library Prep and Sequencing

Extracted DNA was subject to PCR to attach the first sample index. PCR samples were cleaned up using the Zymo DNA Clean and Concentrator kit (Zymo Research, USA) and a second PCR was performed to add a second index and flow cell attachment sequence. PCR products with an expected product size of ∼475 bp were gel-extracted and pooled for Illumina sequencing (Advanced Genomics Core, University of Michigan). Subsequent fastq files were demultiplexed, and target guides were converted into library read counts using an in-house R script modified from the depositor code. All figures and statistical analysis were prepared in R.

### Toxicity experiments

#### Mammalian Cell Culture and Cell Line Maintenance

The HEK-293 and HEPG2 cell lines were purchased from the American Type Culture Collection (ATCC, USA). Cells were cultured and stocks were frozen in Gibco Recovery Cell Culture Freezing Medium at 2 × 10^6^ cells per aliquot. Cell media used for culturing both cell lines was Dulbecco’s Modified Eagle Medium (DMEM, Life Technologies) supplemented with 10% Fetal Bovine Serum (FBS) and 1% penicillin/streptomycin solution (Pen/Strep). Cell flasks were grown at 37°C with 5% CO2.

#### Cell Viability Assays

HEK293 and HEPG2 cells were tested for viability with the fasudil – mecillinam combination in opaque 384-well tissue culture treated plates seeded at 1,000 cells/well. Cells were adhered to the plate overnight and after 24 hours, treated with serial dilutions of fasudil, mecillinam, and a checkerboard assay with fasudil – mecillinam as detailed above. Compounds were dispensed using the Echo liquid handler (Center for Chemical Genomics, University of Michigan). Finally, 20μL of fresh medium was added to each well. Plates were incubated as above for 24 hours. Following incubation, 20μL of CellTiter-Glo reagent (Promega, USA) was added to every well and plates were mixed for 5 minutes on an orbital shaker at room temperature prior to luminescence reading using a spectrophotometer. Read-out was scaled to the DMSO controls and growth curves were plotted using GraphPad Prism 10.4.0.

## DATA AVAILABILITY

Additional information on datasets and results are available in the supplementary materials and the Github Repository (https://github.com/sriram-lab/M2D2). All datasets needed to run the code are provided in the repository. Additional visualizations and heatmap data can be found at https://sriramlab.shinyapps.io/shiny1/.

## CODE AVAILABILITY

All code used within this work is provided through the M2D2 GitHub repository (https://github.com/sriram-lab/M2D2). Additional information on datasets and results are available in the supplementary materials and the Github Repository.

## ACKNOWLEDGEMENTS

We thank the Center for Chemical Genomics for providing high-throughput screening equipment, space and assay design assistance. This project was funded by the Michigan Institute for Data & AI in Society, the Michigan Drug Discovery program, R01AI150826 from National Institute of Allergy and Infectious Diseases, R35GM137795 from National Institute of General Medical Sciences, UM Endowment for Basic Sciences Accelerator Award, and UM Bold Initiatives Research Scouts Award. Main text figures 1-5 were all partially created in BioRender. Reuter, M. (2024) https://BioRender.com/p58p562.

## REFERENCES

1. Centers for Disease Control and Prevention (U.S.). Antibiotic Resistance Threats in the United States, 2019. https://stacks.cdc.gov/view/cdc/82532 (2019) xdoi:10.15620/cdc:82532.

2. Roemhild, R., Bollenbach, T. & Andersson, D. I. The physiology and genetics of bacterial responses to antibiotic combinations. Nat. Rev. Microbiol. (2022) doi:10.1038/s41579-022-00700-5.

3. Tyers, M. & Wright, G. D. Drug combinations: a strategy to extend the life of antibiotics in the 21st century. Nat. Rev. Microbiol. 17, 141–155 (2019).

4. Homer, J. A., Johnson, R. M., Koelln, R. A., Moorhouse, A. D. & Moses, J. E. Strategic re-engineering of antibiotics. Nat Rev Bioeng 1–17 (2024).

5. Challis, G. L. & Hopwood, D. A. Synergy and contingency as driving forces for the evolution of multiple secondary metabolite production by Streptomyces species. Proc. Natl. Acad. Sci. U. S. A. 100 Suppl 2, 14555–14561 (2003).

6. Cokol, M., Li, C. & Chandrasekaran, S. Chemogenomic model identifies synergistic drug combinations robust to the pathogen microenvironment. PLoS Comput. Biol. 14, e1006677 (2018).

7. Cicchese, J. M., Sambarey, A., Kirschner, D., Linderman, J. J. & Chandrasekaran, S. A multi-scale pipeline linking drug transcriptomics with pharmacokinetics predicts in vivo interactions of tuberculosis drugs. Sci. Rep. 11, 5643 (2021).

8. Wildenhain, J. et al. Prediction of Synergism from Chemical-Genetic Interactions by Machine Learning. Cell Syst 1, 383–395 (2015).

9. Jin, W. et al. Deep learning identifies synergistic drug combinations for treating COVID-19. Proc. Natl. Acad. Sci. U. S. A. 118, (2021).

10. Chandrasekaran, S. et al. Chemogenomics and orthology‐based design of antibiotic combination therapies. Mol. Syst. Biol. 12, 872 (2016).

11. Nichols, R. J. et al. Phenotypic landscape of a bacterial cell. Cell 144, 143–156 (2011).

12. Campos, A. I. & Zampieri, M. Metabolomics-Driven Exploration of the Chemical Drug Space to Predict Combination Antimicrobial Therapies. Mol. Cell 74, 1291-1303.e6 (2019).

13. O’Rourke, A. et al. Mechanism-of-Action Classification of Antibiotics by Global Transcriptome Profiling. Antimicrob. Agents Chemother. 64, (2020).

14. Huang, K. et al. DeepPurpose: a deep learning library for drug–target interaction prediction. Bioinformatics 1–6 (2020).

15. Kuwahara, H. & Gao, X. Analysis of the effects of related fingerprints on molecular similarity using an eigenvalue entropy approach. J. Cheminform. 13, 27 (2021).

16. Chou, K.-C. Pseudo Amino Acid Composition and its Applications in Bioinformatics, Proteomics and System Biology. Curr. Proteomics 6, 262–274 (2009).

17. Gilson, M. K. et al. BindingDB in 2015: A public database for medicinal chemistry, computational chemistry and systems pharmacology. Nucleic Acids Res. 44, D1045–53 (2016).

18. Brochado, A. R. et al. Species-specific activity of antibacterial drug combinations. Nature 559, 259–263 (2018).

19. Katzir, I., Cokol, M., Aldridge, B. B. & Alon, U. Prediction of ultra-high-order antibiotic combinations based on pairwise interactions. PLoS Comput. Biol. 15, e1006774 (2019).

20. Russ, D. & Kishony, R. Additivity of inhibitory effects in multidrug combinations. Nat Microbiol 3, 1339– 1345 (2018).

21. Ma, S. et al. Transcriptomic signatures predict regulators of drug synergy and clinical regimen efficacy against tuberculosis. MBio 10, 1–16 (2019).

22. Kanehisa, M. & Goto, S. KEGG: kyoto encyclopedia of genes and genomes. Nucleic Acids Res. 28, 27–30 (2000).

23. Wishart, D. S. et al. DrugBank 5.0: a major update to the DrugBank database for 2018. Nucleic Acids Res. 46, D1074–D1082 (2018).

24. UniProt Consortium. UniProt: the universal protein knowledgebase in 2021. Nucleic Acids Res. 49, D480– D489 (2021).

25. Keseler, I. M. et al. The EcoCyc Database in 2021. Front. Microbiol. 12, 711077 (2021).

26. Adeshina, Y. O., Deeds, E. J. & Karanicolas, J. Machine learning classification can reduce false positives in structure-based virtual screening. Proc. Natl. Acad. Sci. U. S. A. 117, 18477–18488 (2020).

27. Najm, M., Azencott, C.-A., Playe, B. & Stoven, V. Drug target identification with machine learning: How to choose negative examples. Int. J. Mol. Sci. 22, 5118 (2021).

28. Fiscon, G., Conte, F., Farina, L. & Paci, P. SAveRUNNER: A network-based algorithm for drug repurposing and its application to COVID-19. PLoS Comput. Biol. 17, e1008686 (2021).

29. Corsello, S. M. et al. The Drug Repurposing Hub: a next-generation drug library and information resource. Nat. Med. 23, 405–408 (2017).

30. Herrmann, J. Vascular toxic effects of cancer therapies. Nat. Rev. Cardiol. 17, 503–522 (2020).

31. Glass, C. K. & Mitchell, R. N. Winning the battle, but losing the war: mechanisms and morphology of cancer-therapy-associated cardiovascular toxicity. Cardiovasc. Pathol. 30, 55–63 (2017).

32. Basak, D., Arrighi, S., Darwiche, Y. & Deb, S. Comparison of anticancer drug toxicities: Paradigm shift in adverse effect profile. Life (Basel) 12, 48 (2021).

33. Yazbeck, V. et al. An overview of chemotoxicity and radiation toxicity in cancer therapy. Adv. Cancer Res. 155, 1–27 (2022).

34. Shiri, S. et al. Surface Patterned Omniphobic Tiles (SPOTs): a versatile platform for scalable liquid handling. bioRxiv 2024.01.17.575712 (2024) doi:10.1101/2024.01.17.575712.

35. Albo, J. et al. EZ-SPOTs: A simple and robust high-throughput liquid handling platform. bioRxivorg 2024.05.13.594031 (2024).

36. Maier, L. et al. Extensive impact of non-antibiotic drugs on human gut bacteria. Nature 555, 623–628 (2018).

37. Rácz, B. & Spengler, G. Repurposing Antidepressants and Phenothiazine Antipsychotics as Efflux Pump Inhibitors in Cancer and Infectious Diseases. Antibiotics (Basel) 12, (2023).

38. Caldara, M. & Marmiroli, N. Antimicrobial Properties of Antidepressants and Antipsychotics-Possibilities and Implications. Pharmaceuticals 14, (2021).

39. Cui, Z.-H. et al. Phentolamine Significantly Enhances Macrolide Antibiotic Antibacterial Activity against MDR Gram-Negative Bacteria. Antibiotics (Basel) 12, (2023).

40. Romano, K. P. et al. Perturbation-specific transcriptional mapping for unbiased target elucidation of antibiotics. Proc. Natl. Acad. Sci. U. S. A. 121, e2409747121 (2024).

41. Ambrose, P. G. et al. Norepinephrine in combination with antibiotic therapy increases both the bacterial replication rate and bactericidal activity. Antimicrob. Agents Chemother. 62, (2018).

42. Glajzner, P., Bernat, A. & Jasińska-Stroschein, M. Improving the treatment of bacterial infections caused by multidrug-resistant bacteria through drug repositioning. Front. Pharmacol. 15, 1397602 (2024).

43. Silver, L. L. Fosfomycin: Mechanism and resistance. Cold Spring Harb. Perspect. Med. 7, a025262 (2017).

44. Falagas, M. E., Vouloumanou, E. K., Samonis, G. & Vardakas, K. Z. Fosfomycin. Clin. Microbiol. Rev. 29, 321–347 (2016).

45. Thulin, E., Sundqvist, M. & Andersson, D. I. Amdinocillin (Mecillinam) Resistance Mutations in Clinical Isolates and Laboratory-Selected Mutants of Escherichia coli. Antimicrob Agents Chemother 59, 1718– 1727 (2015).

46. Thulin, E. & Andersson, D. I. Upregulation of PBP1B and LpoB in cysB mutants confers mecillinam (amdinocillin) resistance in Escherichia coli. Antimicrob. Agents Chemother. 63, (2019).

47. Typas, A., Banzhaf, M., Gross, C. A. & Vollmer, W. From the regulation of peptidoglycan synthesis to bacterial growth and morphology. Nat. Rev. Microbiol. 10, 123–136 (2011).

48. Chung, C. H. et al. Transfer learning predicts species-specific drug interactions in emerging pathogens. bioRxivorg 2024.06.04.597386 (2024).

49. Hirakawa, H., Suzue, K. & Tomita, H. Roles of the Tol/pal system in bacterial pathogenesis and its application to antibacterial therapy. Vaccines (Basel) 10, 422 (2022).

50. Ozvoldik, K., Stockner, T. & Krieger, E. YASARA Model-interactive molecular modeling from two dimensions to virtual realities. J. Chem. Inf. Model. 63, 6177–6182 (2023).

51. D.A. Case, V. Babin, J.T. Berryman, R.M. Betz, Q. Cai, D.S. Cerutti, T.E. Cheatham, III, T.A. Darden, R.E. Duke, H. Gohlke, A.W. Goetz, S. Gusarov, N. Homeyer, P. Janowski, J. Kaus, I. Kolossváry, A. Kovalenko, T.S. Lee, S. LeGrand, T. Luchko, R. Luo, B. Madej, K.M. Merz, F. Paesani, D.R. Roe, A. Roitberg, C. Sagui, R. Salomon-Ferrer, G. Seabra, C.L. Simmerling, W. Smith, J. Swails, R.C. Walker, J. Wang, R.M. Wolf, X. Wuand P.A. Kollman. AMBER 14. Preprint at (2014).

52. The MathWorks Inc. MATLAB Version: 9.11.0 (R2021b). (2021).

