## Supplemental Materials for "Ultra-high-throughput screening of antimicrobial combination therapies using a two-stage transparent machine learning model"

### SUPPLEMENTARY FIGURES

**A.**

| BINDINGDB | MACCS PseudoAAC | | | |
|---|---|---|---|---|
| Threshold | Accuracy | F1score | Recall | Precision |
| 0 | 0.1845 | 0.3116 | 1 | 0.1845 |
| 0.1 | 0.8402 | 0.6854 | 0.9465 | 0.5381 |
| 0.2 | 0.9131 | 0.7914 | 0.8929 | 0.7107 |
| **0.3** | **0.9315** | **0.819** | **0.8425** | **0.7976** |
| **0.4** | **0.9359** | **0.8224** | **0.7957** | **0.8475** |
| 0.5 | 0.9352 | 0.8108 | 0.7492 | 0.882 |
| 0.6 | 0.9321 | 0.7941 | 0.7036 | 0.9076 |
| 0.7 | 0.9267 | 0.7656 | 0.6509 | 0.9312 |
| 0.8 | 0.9182 | 0.7249 | 0.586 | 0.9525 |
| 0.9 | 0.9056 | 0.6621 | 0.5035 | 0.9714 |
| 1 | 0.8701 | 0.4551 | 0.2978 | 0.9938 |

**B.**

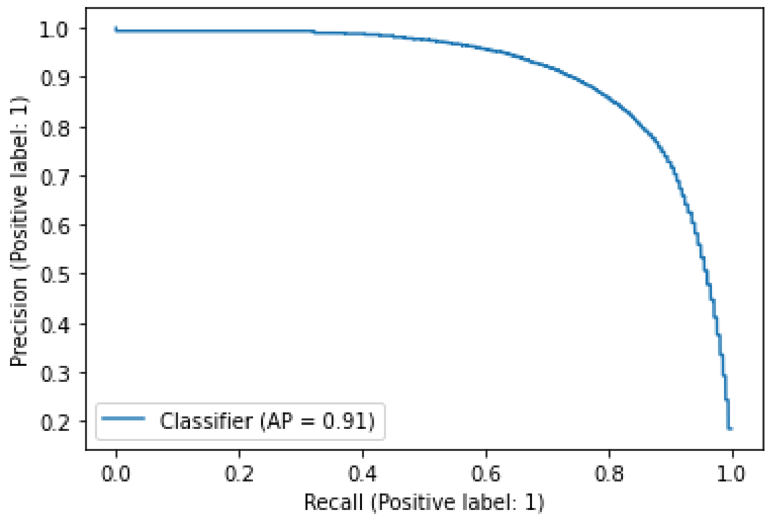

**C.**

| ML model | MSE |
|---|---|
| Neural Network | 1.472 |
| Random Forest | 0.838 |
| Attention based | 1.817 |

**D.**

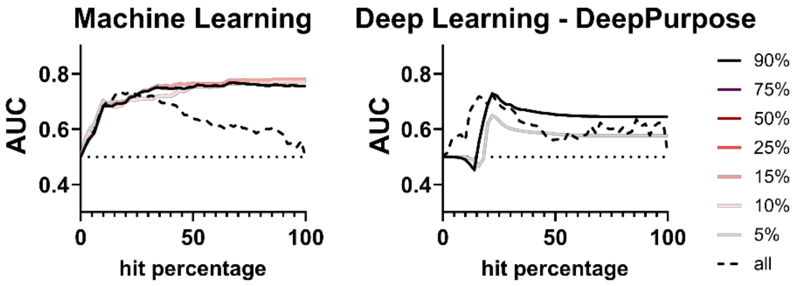

**Supplementary Figure 1.** Validation of the first ML step in M2D2. (A, B) Accuracy, F1 score, recall, precision, and AUC using the BindingDB database as a training set and MACCS keys/PseudoAAC encodings for drugs/proteins. These are the results of a random forest model, which was the best performing algorithm. (C) Mean square error for algorithms tested. The error was calculated from the task of predicting log $K_d$ dissociation constants. (D) The ability of drug – protein interactions from M2D2 vs. DeepPurpose[1] to predict known drug targets. AUC calculation representing a dataset's ability to hit known drug targets was performed to assess the robustness of the calculation over a range of thresholds used to determine the criterion for a strong interaction between a drug and target. An importance filter was added to the calculation to see if the M2D2 model could improve the AUC calculation. The darker the line the more targets were included in the AUC calculation (e.g., 10% uses only the top 10% most important proteins according to ML). The filter was applied from 0 - 100% in increments of 5%. Not every filter is shown for visual clarity.

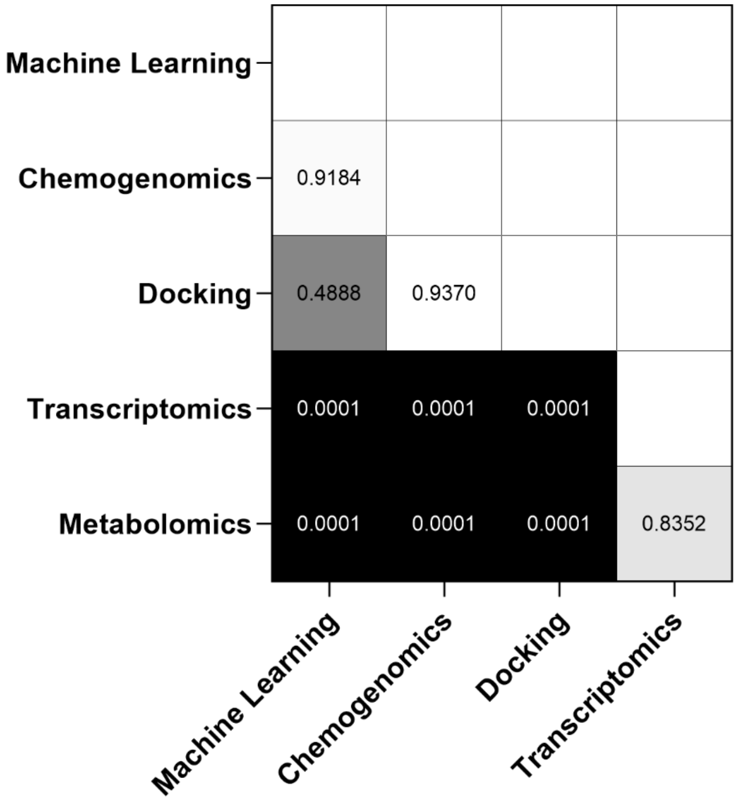

**Supplementary Figure 2.** Adjusted p-value measuring the difference between datasets Pearson's r correlation for 70/30 holdout from main figure text Figure 1. Black indicates significance. Grey indicates no significance. 0.0001 on this figure represents a p-value < 0.0001. Calculations were performed in GraphPad Prism 10.4.0. Pearson's r correlation calculated using 50 iterations of 70/30 hold out method. The graph is sorted from highest to lowest Pearson's r and meaningful differences are noted above the bars using brackets. Three datasets (machine learning, chemogenomics, and molecular docking) have significantly higher correlations compared to the other two datasets (transcriptomics and metabolomics).

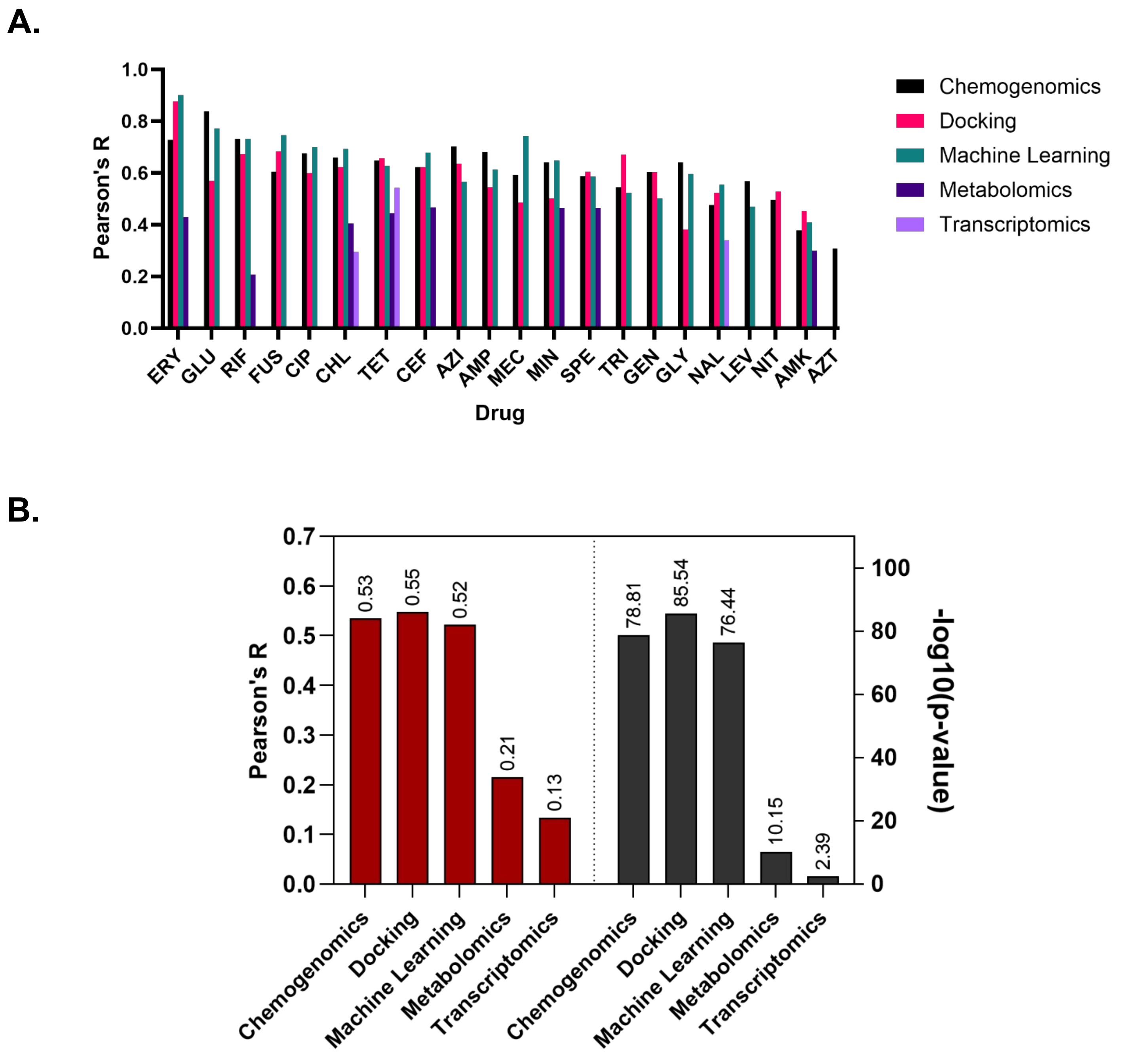

**Supplementary Figure 3.** Leave one out analysis for all possible drugs across datasets. Analysis was done using unweighted and weighted ML with lower weight for high throughput drug interaction datasets (see Supplementary Figure 4). Pictured are the best scores, which were from high-throughput training data weighted 75% as important as non-high throughput analysis. Drugs with less than 25 total training data interactions were excluded. (A) All Pearson's r correlations for each drug left out for each dataset. Pearson's r was calculated using a hold-out method. In this case, all interactions involving a specific drug were withheld from the training set and used as a test set to calculate Pearson's r. Metabolomics and transcriptomics datasets contained fewer drugs, thus some scores are absent for those datasets. R correlations that had p-values that were greater than 0.05 were excluded from the figure. The drugs were sorted by highest average score across chemogenomics, docking, and machine learning. (B) Overall Pearson's r (red) and p-values (black) for the leave one out analysis.

A.

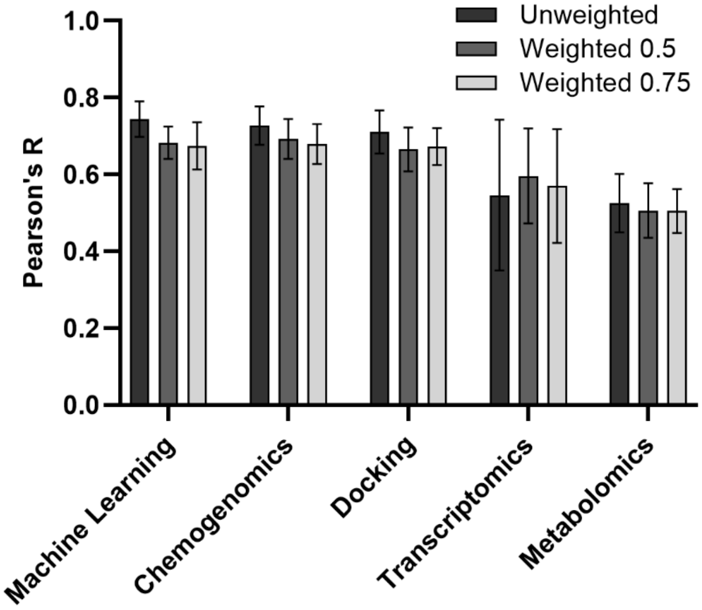

| Dataset | p-value of Unweighted vs. Weighted 0.5 |
|---|---|
| Machine Learning | <0.000001 |
| Chemogenomics | 0.000954 |
| Docking | 0.000107 |
| Transcriptomics | 0.131745 |
| Metabolomics | 0.198539 |

| Dataset | p-value of Unweighted vs. Weighted 0.75 |
|---|---|
| Machine Learning | <0.000001 |
| Chemogenomics | 0.000010 |
| Docking | 0.000373 |
| Transcriptomics | 0.498190 |
| Metabolomics | 0.133836 |

B.

| Dataset | Unweighted | | Weighted 0.5 | | Weighted 0.75 | |
|---|---|---|---|---|---|---|
| | R | pval | R | pval | R | pval |
| MACHINE LEARNING | 0.525907 | 1.40E-76 | 0.536459 | 3.48E-80 | 0.534587 | 1.55E-79 |
| CHEMOGENOMICS | 0.536083 | 4.04E-82 | 0.545302 | 1.86E-85 | 0.547496 | 2.89E-86 |
| DOCKING | 0.422925 | 1.57E-48 | 0.39768 | 1.29E-42 | 0.521833 | 3.66E-77 |
| TRANSCRIPTOMICS | 0.209837 | 1.96E-10 | 0.20191 | 9.43E-10 | 0.214859 | 7.02E-11 |
| METABOLOMICS | 0.141101 | 0.002528 | 0.129533 | 0.005603 | 0.134148 | 0.004108 |

**Supplementary Figure 4.** Evaluation of using weighted versus unweighted training data. Weights were applied to datasets that used high-throughput data collection methods. These high-throughput methods were weighted half or three quarters as much as the low throughput methods. (A) Pearson's r correlation for drug – drug predictions using weighted versus unweighted ML. P-values for t-tests comparing weighted to unweighted are in tables below the figure. (B) Overall Pearson's r correlation and p-values for leave one out drug analysis using weighted or unweighted training data.

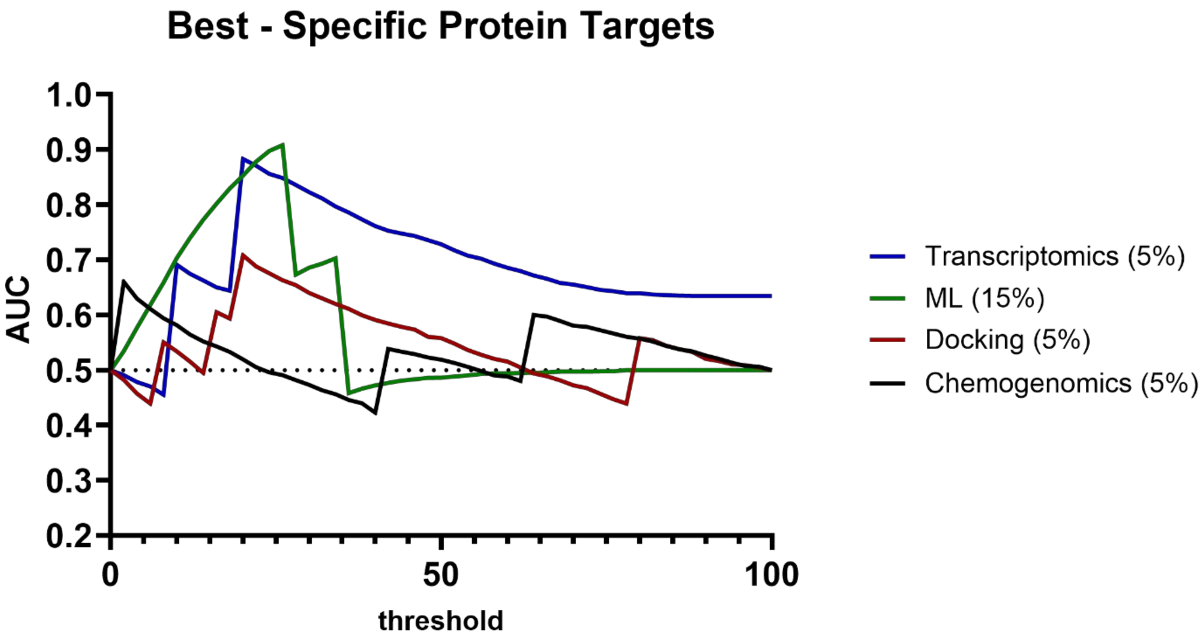

**Supplementary Figure 5.** AUC calculation representing a dataset's ability to hit known drug targets was performed to assess the robustness of the calculation over a range of thresholds. Thresholds in this case indicate the cutoff for a strong interaction between a drug and target. An importance filter was added to the calculation to see if the M2D2 model could improve the AUC calculation. The higher the % more targets were included in the AUC calculation (e.g., 10% uses only the top 10% most important proteins according to ML). The filter was applied from 0 - 100% in increments of 5%. Not every filter is shown for visual clarity. The best M2D2 ML importance filter was chosen and represented in the graph with the importance percentage in parentheses. Specific protein targets were taken from DrugBank[2].

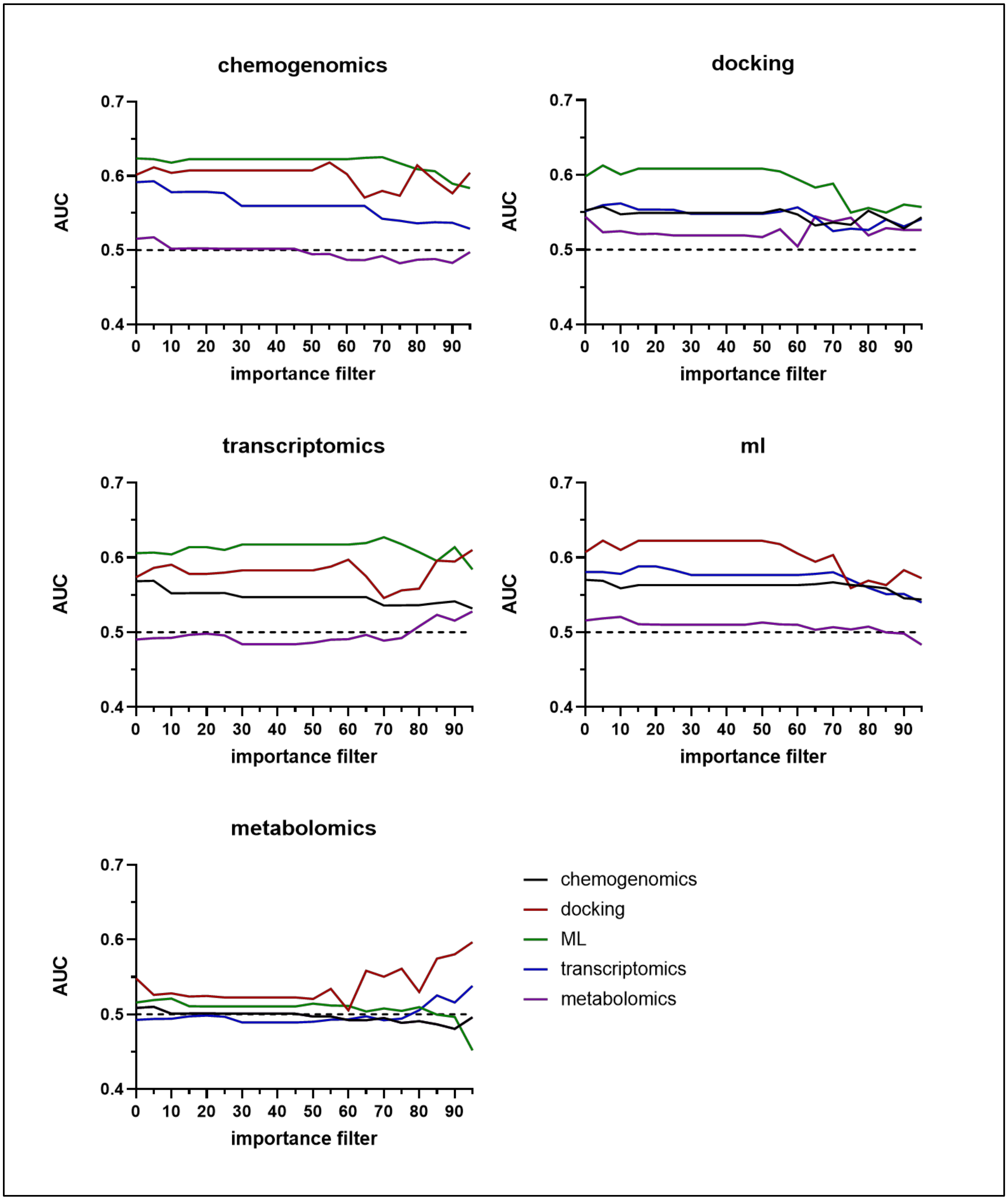

**Supplementary Figure 6.** AUC calculation representing a dataset's ability to match another dataset. These calculations were all performed where recall of primary Kegg targets was 0.50. The title of the graph indicates the dataset being matched to. An importance filter was added to the calculation to see if the M2D2 model could improve the AUC calculation. The higher the filter the fewer targets were included in the AUC calculation (e.g., 10 uses only the top 90% most important proteins according to ML).

**A.**

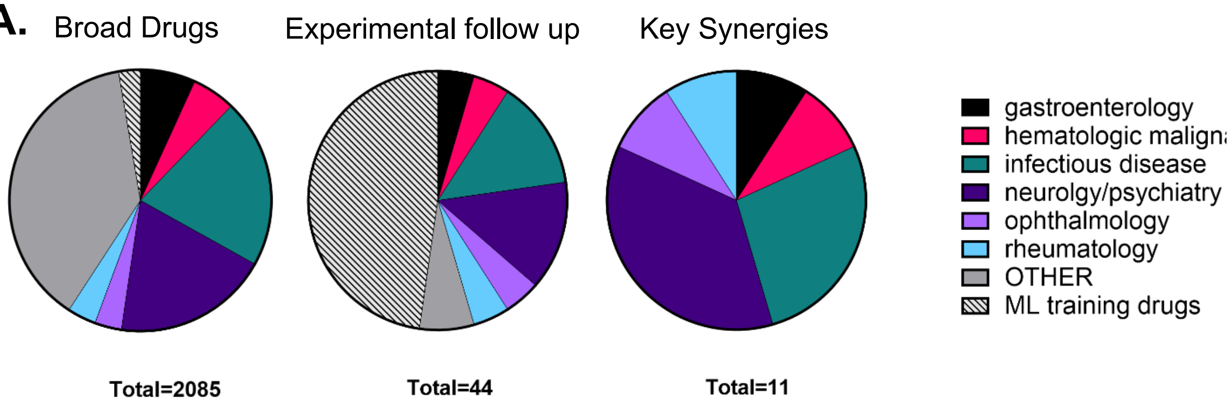

**B.**

| full_name | full_name_Broad | Code ML | Code Exp |
|---|---|---|---|
| AG-120 (ivosidenib) | ivosidenib | ivosidenib | AG |
| amoxicillin | amoxicillin | AMX | AMO |
| ampicillin | ampicillin | AMP | AMP |
| azithromycin | azithromycin | AZI | AZI |
| bortezomib | bortezomib | bortezomib | BOR |
| cefaclor | cefaclor | CEC | CEFA |
| cefsulodin | cefsulodin | CFS | CEFS |
| chloramphenicol | chloramphenicol | CHL | CHL |
| chlorpropamide | chlorpropamide | chlorpropamide | CHPROP |
| ciprofloxacin | ciprofloxacin | CIP | CIP |
| clarithromycin | clarithromycin | CLA | CLA |
| clopamide | clopamide | clopamide | CLO |
| dolasetron | dolasetron | dolasetron | DOL |
| erythromycin | erythromycin | ERY | ERY |
| etomidate | etomidate | etomidate | ETM |
| etoricoxib | etoricoxib | etoricoxib | ETX |
| fluphenazine | fluphenazine | fluphenazine | FLU |
| fosamprenavir | fosamprenavir | fosamprenavir | FOS |
| fusidic acid | fusidic-acid | FUS | FUS |
| HA-1077 (fasudil) | fasudil | fasudil | HA |
| mecillinam | pivmecillinam | MEC | MEC |
| nimesulide | nimesulide | nimesulide | NIM |
| nitrofurantoin | nitrofurantoin | NIT | NIT |
| polymyxin b | polymyxin-B-sulfate | PMB | PMB |
| PTC-124 (ataluren) | ataluren | ataluren | PTC |
| puromycin | puromycin | PUR | PUR |
| rifampicin | rifampin | RIF | RIF |
| ripasudil | ripasudil | ripasudil | RIP |
| simeprevir | simeprevir | simeprevir | SIM |
| succinylcholine-chloride | succinylcholine-chloride | succinylcholine-chloride | SUC |
| sulfachlorpyridazine | sulfachlorpyridazine | sulfachlorpyridazine | SCP |
| sulfadiazine | sulfadiazine | sulfadiazine | SUD |
| sulfamerazine | sulfamerazine | sulfamerazine | SMR |
| sulfamethazine | sulfamethazine | sulfamethazine | SMT |
| sulfamethoxazole | sulfamethoxazole | SMX | SMZ |
| sulfaquinoxaline | sulfaquinoxaline | sulfaquinoxaline | SULQ |
| talniflumate | talniflumate | talniflumate | TAL |
| tetracycline | tetracycline | TET | TET |
| tobramycin | tobramycin | TOB | TOB |
| toltrazuril | toltrazuril | toltrazuril | TOL |
| trimethoprim | trimethoprim | TRI | TMP |
| vancomycin | vancomycin | VAN | VAN |
| verapamil | verapamil | VER | VER |
| ziprasidone | ziprasidone | ziprasidone | ZIP |

**Supplementary Figure 7.** Overview of repurposed drugs chosen for computational and experimental work. (A) Breakdown of drug categories for the set of repurposed drugs used for M2D2 predictions, experimental high-throughput screening, and key synergistic pairs predicted via M2D2 and confirmed with high-throughput experiments. (B) List of drug names and abbreviations used for high-throughput experiments. Full name, Full name from the Broad Institute repurposing hub, abbreviation for ML calculations, and abbreviation for high-throughput experiments are noted.

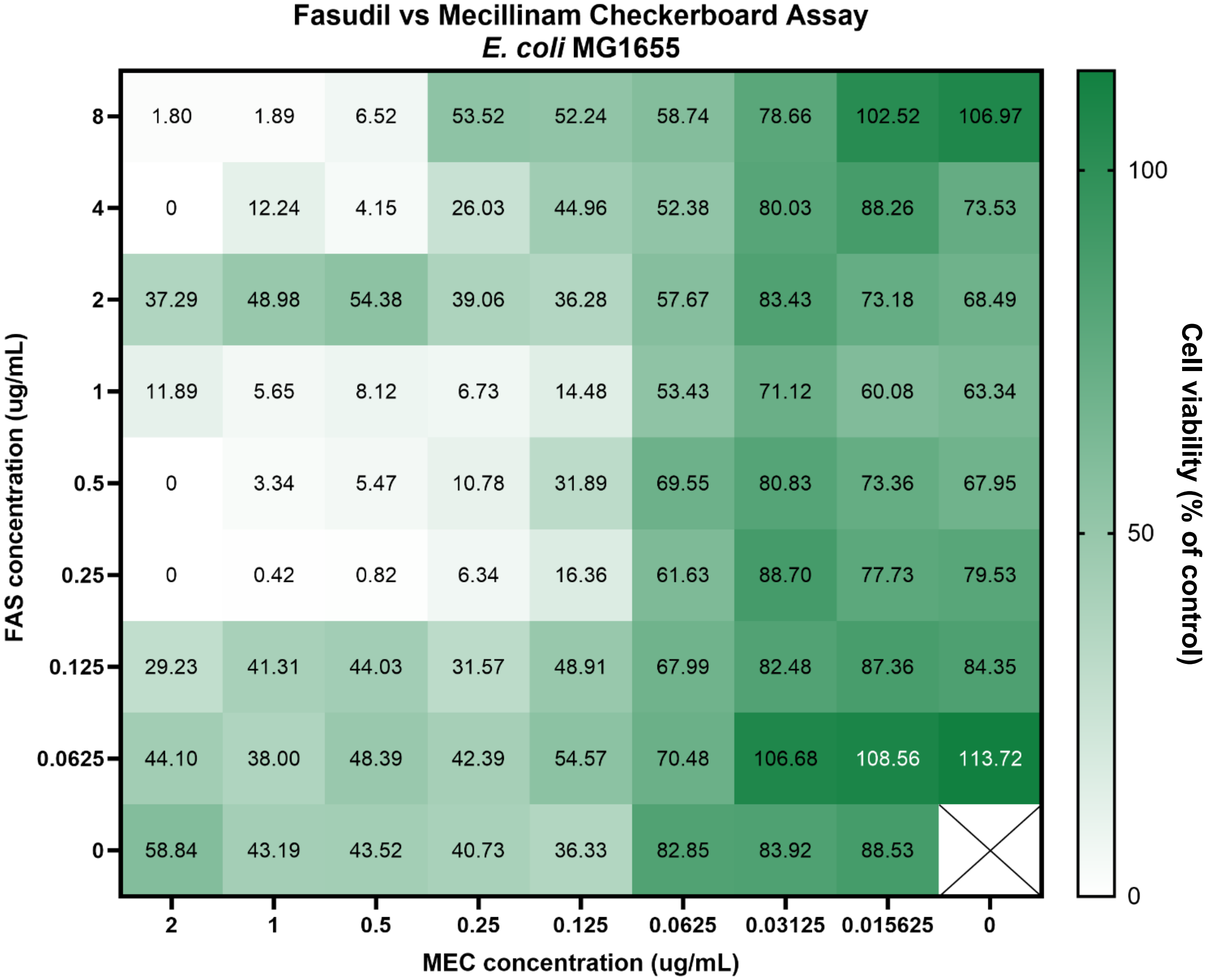

**Supplementary Figure 8**. Full traditional checkerboard assay in *E. coli* MG1655 with mecillinam and fasudil. Each compound was serially diluted 2-fold at a starting concentration at 2 x minimum inhibitory concentration for each compound. These values are the mean of three replicates.

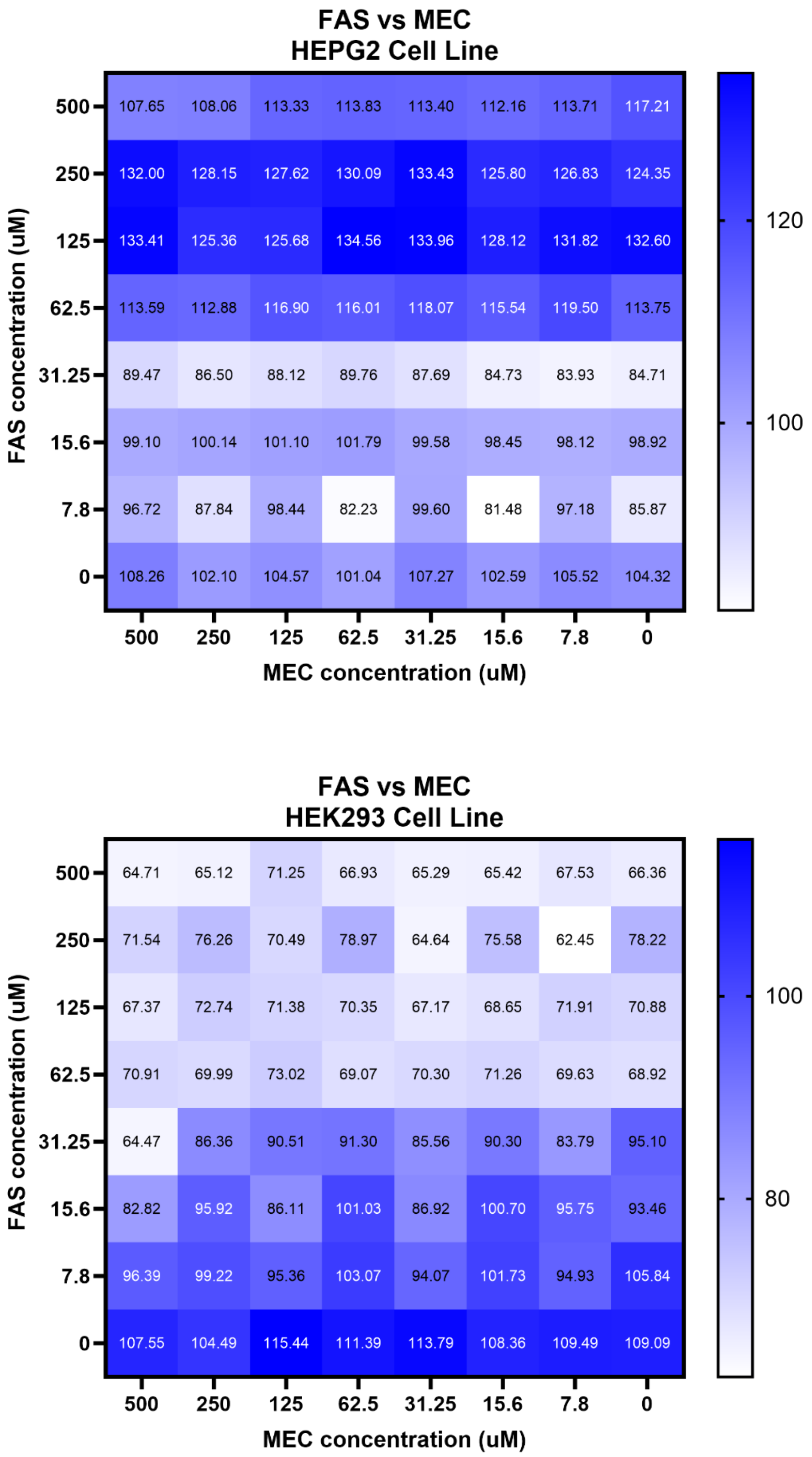

**Supplementary Figure 9.** Toxicity experiments were performed for human liver (HEPG2) and kidney (HEK293) cell lines. Growth curves are shown in the heatmap. The HEK293 figure is the full toxicity assay, shown in Figure 5c of the main text. The darker the blue the less toxic the combination to liver or kidney cells.

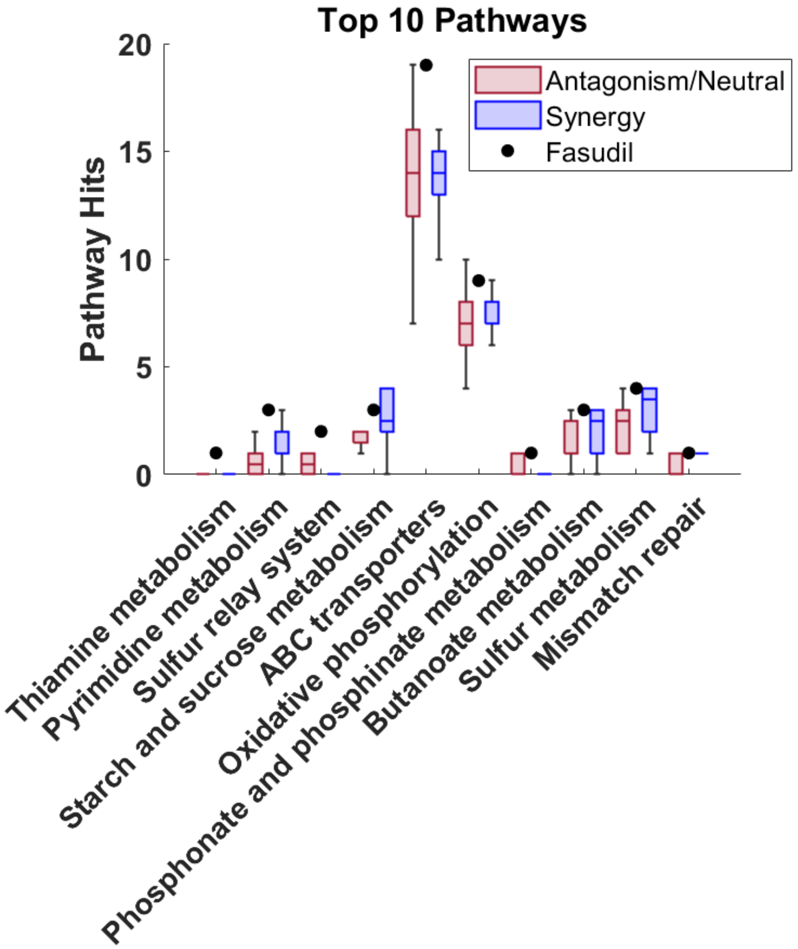

**Supplementary Figure 10.** The top 10 pathways predicted to be hit by fasudil by M2D2 (percentile cutoff 80, importance filter of 15%). Pathways were sorted and ranked based on p-value of t-test between fasudil and the group of drugs that have an antagonistic or neutral interaction with mecillinam. Fasudil shows higher hits in Sulfur metabolism and pyrimidine metabolism, compared to drugs that are antagonistic with mecillinam, and shows similarity to drugs that are synergistic with mecillinam (blue boxes).

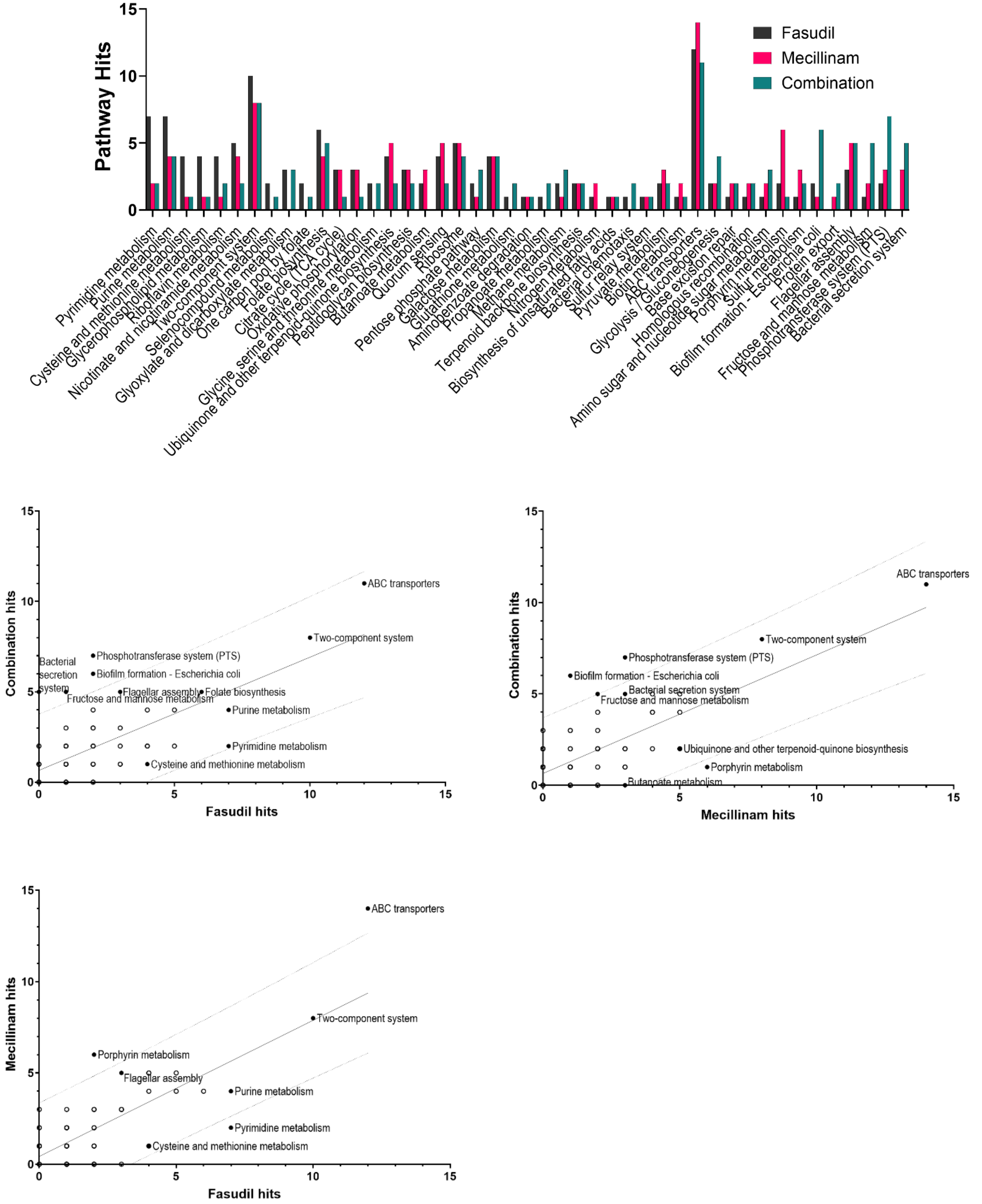

**Supplementary Figure 11.** CRISPRi pathway hits for fasudil, mecillinam, and their combination represented in bar and scatter plots. The bar graph is sorted by the difference in fasudil hits and average of mecillinam and combination hits. Scatter plots show fasudil vs. combination, mecillinam vs. combination, and fasudil vs. mecillinam. Pathways that are outside the 95% confidence prediction bands are labeled, as are pathways with many hits from either dataset. Correlation and prediction bands were done in GraphPad Prism 10.4.0.

### SUPPLEMENTARY TABLES

**A.**

| Drug | R | -LOG10 (p-values) | Drug size (heavy atoms) | Target | Static, Cidal, Both |
|---|---|---|---|---|---|
| ERY | 0.900 | 11.63 | **51** | Protein synthesis, 50S | S |
| GLU | 0.771 | 5.81 | **12** | NA | |
| FUS | 0.747 | 5.67 | **37** | Elongation factor—protein synthesis | S |
| MEC | 0.743 | 4.68 | **22** | Cell wall | C |
| RIF | 0.731 | 15.69 | **59** | RNA synthesis | C |
| CIP | 0.699 | 8.39 | **24** | DNA gyrase | C |
| CHL | 0.693 | 11.96 | **20** | Protein synthesis, 50S | S |
| CEF | 0.678 | 4.69 | **28** | Cell wall | C |
| MIN | 0.647 | 3.45 | **33** | Protein synthesis, 16S | S |
| TET | 0.627 | 6.19 | **32** | Protein synthesis, 30S | S |
| AMP | 0.613 | 5.86 | **24** | Cell wall | C-S |
| GLY | 0.596 | 5.79 | **6** | NA | |
| SPE | 0.586 | 5.49 | **17** | Protein synthesis, 30S | S |
| AZI | 0.565 | 5.86 | **52** | Protein synthesis, 50S | S |
| NAL | 0.554 | 4.43 | **17** | DNA gyrase | C |
| TRI | 0.523 | 3.27 | **21** | Folic acid biosynthesis | S |
| GEN | 0.501 | 2.04 | **33** | Protein synthesis, 30S | C-S |
| LEV | 0.468 | 1.74 | **26** | DNA gyrase | C |
| AMK | 0.409 | 2.10 | **40** | Protein synthesis, 30S | C-S |
| PMS | 0.341 | 1.12 | **6** | Oxidative stress | C |
| NVB | 0.267 | 0.73 | **44** | DNA gyrase | S |
| AZT | 0.257 | 1.13 | **28** | Cell wall | C |
| NIT | 0.197 | 0.74 | **17** | Multiple, DNA | C-S |
| FOS | 0.185 | 0.45 | **8** | Cell wall | C |
| TRC | 0.097 | 0.22 | **17** | Fatty acid biosynthesis | S |
| PRC | 0.074 | 0.14 | **17** | NA | |

**B.**

| CHEMOGENOMICS | | | DOCKING | | | MACHINE LEARNING | | | METABOLOMICS | | |
|---|---|---|---|---|---|---|---|---|---|---|---|
| Drug | R | p-value | Drug | R | p-value | Drug | R | p-value | Drug | R | p-value |
| GLU | 0.838 | 2.6E-08 | ERY | 0.876509 | 4.87E-11 | ERY | 0.900224 | 2.34E-12 | MIN | 0.662064 | 0.00023 |
| RIF | 0.732 | 1.76E-16 | FUS | 0.682097 | 3.31E-05 | GLU | 0.771205 | 1.56E-06 | CEF | 0.465818 | 0.007212 |
| ERY | 0.727 | 2.47E-06 | RIF | 0.67272 | 2.81E-13 | FUS | 0.746661 | 2.15E-06 | SPE | 0.463385 | 0.000418 |
| AZI | 0.702632 | 1.38E-10 | TRI | 0.671403 | 2.11E-06 | MEC | 0.742847 | 2.11E-05 | TET | 0.444808 | 0.000954 |
| AMP | 0.680448 | 2.87E-08 | TET | 0.656545 | 1.26E-07 | RIF | 0.730658 | 2.03E-16 | ERY | 0.429115 | 0.014256 |

**Supplementary Table 1.** Leave one out analysis for all possible drugs across datasets. Analysis was done using unweighted and weighted ML. Pictured are the best scores, which were from high-throughput training data weighted 75% as important as non-high throughput analysis. Drugs with less than 25 total training data interactions were excluded. (A) Table of all scores for machine learning input. (B) Top 5 R correlation scores for each dataset. Transcriptomics is excluded because only 3 scores had an associated p-value of at least less than 0.05. Drugs that appear in at least three datasets are highlighted in green.

| | | | | | | | | |
|---|---|---|---|---|---|---|---|---|
| | Cutoff 80 | | | | | | | |
| Filter | R_mean | | | | R_std | | | |
| | HH | LL | HL | LH | HH | LL | HL | LH |
| 1 | 0.452 | 0.578 | 0.521 | 0.546 | 0.055 | 0.065 | 0.048 | 0.056 |
| 2 | 0.465 | 0.518 | 0.563 | 0.521 | 0.056 | 0.056 | 0.052 | 0.047 |
| 3 | 0.479 | 0.297 | 0.263 | 0.178 | 0.053 | 0.062 | 0.049 | 0.059 |
| 4 | 0.541 | 0.164 | 0.258 | 0.199 | 0.060 | 0.075 | 0.062 | 0.057 |
| 5 | 0.560 | 0.289 | 0.319 | 0.227 | 0.060 | 0.065 | 0.058 | 0.074 |
| 6 | 0.639 | 0.167 | 0.382 | 0.003 | 0.078 | 0.170 | 0.159 | 0.202 |
| 7 | 0.574 | 0.254 | 0.474 | 0.166 | 0.114 | 0.213 | 0.150 | 0.185 |
| | Cutoff 80 | | | | Cutoff 95 | | | |
| Filter | R_mean | | R_std | | R_mean | | R_std | |
| | mean | median | | | mean | median | | |
| 1 | 0.614 | 0.629 | 0.047 | 0.041 | 0.610 | 0.606 | 0.043 | 0.048 |
| 2 | 0.586 | 0.583 | 0.040 | 0.051 | 0.592 | 0.593 | 0.044 | 0.046 |
| 3 | 0.611 | 0.609 | 0.046 | 0.044 | 0.607 | 0.586 | 0.043 | 0.038 |
| 4 | 0.643 | 0.656 | 0.049 | 0.056 | 0.669 | 0.666 | 0.048 | 0.039 |
| 5 | 0.622 | 0.613 | 0.053 | 0.042 | 0.598 | 0.594 | 0.044 | 0.051 |
| 6 | 0.609 | 0.581 | 0.073 | 0.076 | 0.582 | 0.595 | 0.091 | 0.070 |
| 7 | 0.609 | 0.602 | 0.110 | 0.110 | 0.613 | 0.587 | 0.102 | 0.103 |
| | Cutoff 90 | | | | Cutoff 85 | | | |
| | R_mean | | R_std | | R_mean | | R_std | |
| | mean | median | | | mean | median | | |
| 4 | 0.651 | 0.642 | 0.049 | 0.042 | 0.647 | 0.649 | 0.051 | 0.045 |

**Supplementary Table 2.** Table of Pearson's r and corresponding standard deviation for M2D2 trained on the output of high-throughput experiments using the SPOTs method[3,4]. HH, LL, HL, and LH represent drug 1 and drug 2 doses (H = high dose, L = low dose). High and low concentrations were determined based on MIC20 (high) and MIC80 (low). Measurements were taken at two time points 8 and 16 hours and done in duplicate. To provide higher confidence in the SPOTs predictions, statistical features were applied to the data and was organized in the following structure: (1) time point one, (2) time point 2, (3) standard deviation of each drug – drug combination less than 5000 for time point one, (4) standard deviation of each drug – drug combination less than 5000 for time point two, (5) consistent synergy, antagonism, or neutral scores over both time points, (6) same criteria as 5 and a standard deviation in the single drug MIC was less than 5000, (7) same criteria as 6 and multiple concentrations show the same interaction category (synergy/antagonism/neutral). A range of cutoffs used to determine the criterion for a strong interaction between a drug and target was used to improve the robustness of the analysis. Using the mean or median scores of the four dose combinations yielded better R values, as did the fourth filtering step.

| Drug (concentration) | Summary | Adjusted P-value |
|---|---|---|
| FAS (8µg/mL) vs. MEC (2µg/mL) | ** | 0.0017 |
| FAS (8µg/mL) vs. FAS + MEC (8µg/mL, 2µg/mL) | **** | <0.0001 |
| FAS (8µg/mL) vs. FAS + MEC (0.25µg/mL, 0.125µg/mL) | **** | <0.0001 |
| MEC (2µg/mL) vs. FAS + MEC (8µg/mL, 2µg/mL) | *** | 0.0005 |
| MEC (2µg/mL) vs. FAS + MEC (0.25µg/mL, 0.125µg/mL) | ** | 0.0037 |
| FAS + MEC (8µg/mL, 2µg/mL) vs. FAS + MEC (0.25µg/mL, 0.125µg/mL) | ns | 0.3236 |

**Supplementary Table 3.** Ordinary one-way ANOVA for fasudil (FAS) and mecillinam (MEC) Minimum Inhibitory Concentration (MIC) assays shown in Figure 5 of the main text. Fasudil vs. mecillinam MIC assay was done in E. coli MG1655. P-values were calculated for single drug viability versus combination viability at different concentrations. P-values are from ANOVA Tukey’s multiple comparisons test in GraphPad Prism 10.4.0.

| | cutoff: 75, ML filter: none | | | cutoff: 75, ML filter: top 15% | |
|---|---|---|---|---|---|
| | pathways | p_values | | pathways | p_values |
| 1 | Galactose metabolism | 2.56E-09 | 1 | Thiamine metabolism | 3.62E-08 |
| 2 | Pyrimidine metabolism | 1.36E-07 | 2 | Sulfur relay system | 2.88E-07 |
| 3 | C5-Branched dibasic acid metabolism | 2.19E-06 | 3 | Galactose metabolism | 7.1E-07 |
| 4 | Starch and sucrose metabolism | 2.64E-06 | 4 | Nicotinate and nicotinamide metabolism | 2.75E-06 |
| 5 | Sulfur relay system | 1.32E-05 | 5 | Pyrimidine metabolism | 4.85E-06 |
| 6 | Lipopolysaccharide biosynthesis | 1.76E-05 | 6 | Pyruvate metabolism | 1.32E-05 |
| 7 | Protein export | 4.23E-05 | 7 | Butanoate metabolism | 0.000104 |
| 8 | Purine metabolism | 5.38E-05 | 8 | ABC transporters | 0.000115 |
| 9 | Glutathione metabolism | 6.04E-05 | 9 | Lipopolysaccharide biosynthesis | 0.000193 |
| 10 | Butanoate metabolism | 0.000115 | 10 | Fatty acid biosynthesis | 0.000526 |

| | cutoff: 80, ML filter: none | | | cutoff: 80, ML filter: 15% | |
|---|---|---|---|---|---|
| | pathways | p_values | | pathways | p_values |
| 1 | Folate biosynthesis | 1.09E-08 | 1 | Thiamine metabolism | 1.94E-10 |
| 2 | Phosphonate and phosphinate metabolism | 3.54E-07 | 2 | Pyrimidine metabolism | 1.14E-09 |
| 3 | Sulfur relay system | 1.02E-06 | 3 | Sulfur relay system | 6.71E-09 |
| 4 | Pyrimidine metabolism | 1.34E-06 | 4 | Starch and sucrose metabolism | 2.75E-06 |
| 5 | Starch and sucrose metabolism | 2.53E-06 | 5 | ABC transporters | 1.77E-05 |
| 6 | Sulfur metabolism | 2.99E-06 | 6 | Oxidative phosphorylation | 1.88E-05 |
| 7 | Thiamine metabolism | 7.01E-06 | 7 | Phosphonate and phosphinate metabolism | 3.88E-05 |
| 8 | Biotin metabolism | 7.01E-06 | 8 | Butanoate metabolism | 3.88E-05 |
| 9 | Fructose and mannose metabolism | 1.22E-05 | 9 | Sulfur metabolism | 4.59E-05 |
| 10 | Mismatch repair | 0.000158 | 10 | Mismatch repair | 0.000158 |

| | cutoff: 85, ML filter: none | | | cutoff: 85, ML filter: 15% | |
|---|---|---|---|---|---|
| | pathways | p_values | | pathways | p_values |
| 1 | Glycolysis / Gluconeogenesis | 5.1E-07 | 1 | Glycolysis / Gluconeogenesis | 1.61E-07 |
| 2 | Sulfur metabolism | 8.8E-06 | 2 | Bacterial secretion system | 2.44E-06 |
| 3 | Pyrimidine metabolism | 7.79E-05 | 3 | Glycerophospholipid metabolism | 3.44E-06 |
| 4 | Amino sugar and nucleotide sugar metabolism | 0.000337 | 4 | Sulfur metabolism | 4.28E-06 |
| 5 | Sulfur relay system | 0.000526 | 5 | Sulfur relay system | 3.88E-05 |
| 6 | ABC transporters | 0.00194 | 6 | ABC transporters | 4.31E-05 |
| 7 | Ubiquinone and other terpenoid-quinone biosynthesis | 0.002363 | 7 | Oxidative phosphorylation | 7.08E-05 |
| 8 | O-Antigen nucleotide sugar biosynthesis | 0.002757 | 8 | Pyrimidine metabolism | 0.000158 |
| 9 | Porphyrin metabolism | 0.002757 | 9 | Two-component system | 0.000204 |
| 10 | Flagellar assembly | 0.002819 | 10 | Flagellar assembly | 0.000764 |

| | cutoff: 90, ML filter: none | | | cutoff: 90, ML filter: 15% | |
|---|---|---|---|---|---|
| | pathways | p_values | | pathways | p_values |
| 1 | Amino sugar and nucleotide sugar metabolism | 3.53E-09 | 1 | Amino sugar and nucleotide sugar metabolism | 3.45E-09 |
| 2 | Quorum sensing | 5.98E-06 | 2 | Sulfur metabolism | 1.48E-06 |
| 3 | O-Antigen nucleotide sugar biosynthesis | 7.01E-06 | 3 | Bacterial secretion system | 1.48E-06 |
| 4 | Bacterial secretion system | 1.67E-05 | 4 | Pyruvate metabolism | 7.01E-06 |
| 5 | Glycolysis / Gluconeogenesis | 0.000104 | 5 | ABC transporters | 1.14E-05 |
| 6 | Starch and sucrose metabolism | 0.000158 | 6 | Quorum sensing | 1.34E-05 |
| 7 | Glyoxylate and dicarboxylate metabolism | 0.000158 | 7 | Fructose and mannose metabolism | 2.38E-05 |
| 8 | Nitrogen metabolism | 0.000158 | 8 | Glyoxylate and dicarboxylate metabolism | 0.000158 |
| 9 | Fructose and mannose metabolism | 0.000364 | 9 | Methane metabolism | 0.000158 |
| 10 | Teichoic acid biosynthesis | 0.001272 | 10 | Nitrogen metabolism | 0.000158 |

| | cutoff: 95, ML filter: none | | | cutoff: 95, ML filter: 15% | |
|---|---|---|---|---|---|
| | pathways | p_values | | pathways | p_values |
| 1 | Sulfur metabolism | 8.17E-06 | 1 | Sulfur metabolism | 7.83E-07 |
| 2 | Oxidative phosphorylation | 1.56E-05 | 2 | Quorum sensing | 0.00037 |
| 3 | Ubiquinone and other terpenoid-quinone biosynthesis | 2.54E-05 | 3 | Phosphotransferase system (PTS) | 0.001433 |
| 4 | Lipopolysaccharide biosynthesis | 0.000158 | 4 | Two-component system | 0.002757 |
| 5 | Protein export | 0.000171 | 5 | Ribosome | 0.027068 |
| 6 | Nitrogen metabolism | 0.000526 | | | |
| 7 | ABC transporters | 0.001177 | | | |
| 8 | Bacterial secretion system | 0.001406 | | | |
| 9 | Quorum sensing | 0.001502 | | | |
| 10 | Starch and sucrose metabolism | 0.003372 | | | |

**Supplementary Table 4**. Top ten pathways predicted by M2D2 to be hit at various thresholds, with (blue highlight) and without an M2D2 importance filter (green highlighted). Thresholds determine a strong interaction between drug and protein. 15% importance filter was chosen based on best drug – protein AUC for matching Kegg pathway data. Pathways are highlighted if they appear in the top ten regardless of cutoff. The darker the highlight the more often the pathway appears. None of the green or blue highlighted pathwas appeared in the bottom ten predicted pathways. ABC transporters was a very common pathway but also appeared in the bottom ten predicted pathways for some cutoffs and was therefore highlighted in pink.

| | RMSE | p-value | R |
|---|---|---|---|
| cutoff: 75 ML filter: none | 5.85 | 2.52E-14 | 0.670 |
| cutoff: 80 ML filter: none | 5.06 | 6.85E-14 | 0.661 |
| cutoff: 85 ML filter: none | 4.08 | 1.45E-12 | 0.634 |
| cutoff: 90 ML filter: none | 2.82 | 7.72E-11 | 0.593 |
| cutoff: 95 ML filter: none | 1.29 | 3.27E-09 | 0.549 |

| | RMSE | p-value | R |
|---|---|---|---|
| cutoff: 75 ML filter: 15% | 2.57 | 8.77E-12 | 0.616 |
| cutoff: 80 ML filter: 15% | 2.06 | 3.24E-13 | 0.648 |
| cutoff: 85 ML filter: 15% | 1.34 | 2.49E-12 | 0.629 |
| cutoff: 90 ML filter: 15% | 0.89 | 1.35E-08 | 0.531 |
| cutoff: 95 ML filter: 15% | 0.16 | 9.36E-04 | 0.326 |

| | RMSE | p-value | R |
|---|---|---|---|
| Ampicillin | 5.54 | 3.88E-06 | 0.443 |
| Amoxicillin | 5.56 | 7.61E-06 | 0.431 |
| Gentamicin | 4.60 | 3.22E-08 | 0.519 |
| Mecillinam | 3.69 | 2.70E-22 | 0.787 |
| Sulfamethoxazole | 6.65 | 5.46E-05 | 0.392 |
| Trimethoprim | 5.69 | 1.14E-05 | 0.423 |
| Vancomycin | 4.57 | 8.28E-03 | 0.263 |

**Supplementary Table 5.** The correlations of CRISPRi pathway hits versus M2D2 pathway hits for fasudil over a range of cutoffs that determine a significant interaction between drug and protein in M2D2. The second table also applies an important features filter from the M2D2 model. The third table shows the correlations of hits accorss a range of drugs, including mecillinam, at an M2D2 threshold cuttoff of 80.

| UniProt ID | Protein Full Name (UniProt) | Associated Pathways (Kegg) | ML Score | CRISPRi Log Fold Change |
|---|---|---|---|---|
| P77736 | Putative ankyrin repeat protein YahD | | 0.42 | -2.81 |
| Q47539 | Taurine transport system permease protein TauC | Sulfur metabolism, ABC transporters | 0.48 | -2.81 |
| P0AAX6 | Uncharacterized protein McbA (MqsR-controlled colanic acid and biofilm protein A) | | 0.41 | -2.81 |
| P0AAZ0 | Inner membrane protein YbjO | | 0.38 | -2.81 |
| P75917 | Uncharacterized protein YmdA | | 0.39 | -2.18 |
| P0ABX2 | Flagellar basal-body rod protein FlgC (Putative proximal rod protein) | Flagellar assembly | 0.38 | -3.81 |
| P75968 | Uncharacterized protein YmfE | | 0.35 | -2.81 |
| P11350 | Respiratory nitrate reductase 1 gamma chain (EC 1.7.5.1) (Cytochrome B-NR) (Nitrate reductase A subunit gamma) (Quinol-nitrate oxidoreductase subunit gamma) | Nitrogen metabolism, Two-component system | 0.41 | -3.16 |
| P0AFH6 | Oligopeptide transport system permease protein OppC | Quorum sensing, ABC transporters | 0.37 | -3.61 |
| P0ACW4 | Uncharacterized protein YdcA | | 0.43 | -2.81 |
| P76236 | Probable diguanylate cyclase CdgI (DGC) (EC 2.7.7.65) | | 0.48 | -3.07 |
| P69805 | PTS system mannose-specific EIID component (EII-M-Man) (EIID-Man) (Mannose permease IID component) | Phosphotransferase system (PTS), Fructose and mannose metabolism, Amino sugar and nucleotide sugar metabolism | 0.43 | -2.34 |
| P76264 | Probable manganese efflux pump MntP | | 0.51 | -2.81 |
| P33219 | Protein YebF | | 0.37 | -2.39 |
| P36561 | Adenosylcobinamide-GDP ribazoletransferase (EC 2.7.8.26) (Cobalamin synthase) (Cobalamin-5'-phosphate synthase) | Porphyrin metabolism | 0.36 | -2.41 |
| P33015 | UPF0394 inner membrane protein YeeE | | 0.41 | -2.58 |
| P64536 | Uncharacterized protein YeiS | | 0.41 | -2.81 |
| P33021 | Putative nucleoside permease NupX | | 0.39 | -2.81 |
| P0ABW3 | Uncharacterized ferredoxin-like protein YfaE | | 0.42 | -2.81 |
| P32139 | Uncharacterized protein YihR | Galactose metabolism, Glycolysis / Gluconeogenesis | 0.41 | -2.61 |
| P0A9E2 | Regulatory protein SoxS | | 0.42 | -2.63 |
| P0AFJ1 | Protein YjdM | | 0.36 | -2.59 |
| P39282 | Inner membrane transporter YjeM | | 0.50 | -2.81 |
| P0ADD9 | Uncharacterized protein YjjY | | 0.42 | -3.81 |
| P0ADB4 | Entericidin A | | 0.44 | -3.81 |
| Q2EET2 | UPF0370 protein YpfN | | 0.44 | -2.81 |
| A5A628 | Uncharacterized protein YjbT | | 0.38 | -2.81 |
| P58036 | Uncharacterized protein YjbS | | 0.45 | -2.81 |

**Supplementary Table 6.** A list of proteins that overlap between M2D2 drug – protein interactions and CRISPRi negative log fold change genes. More specifically, this list is the overlap between hits by the drug fasudil (top 80$^{th}$ percentile of interaction scores) predicted by M2D2 and genes that, when knocked down by CRISPRi, resulted in loss of fitness. The M2D2 importance filter of 15% was also applied to the list of proteins. Full names of the proteins were taken from UniProt[5]. If available, the Kegg database[6] pathways that the protein is a part of are noted. The list is sorted by the largest total score between ML and the absolute value of the CRISPRi log fold change.

| UniProt ID | Protein Full Name (UniProt) | Associated Pathways (Kegg) | ML Score | CRISPRi Log Fold Change |
|---|---|---|---|---|
| P0AAA1 | Inner membrane protein YagU | | 0.43 | -3.81 |
| P77736 | Putative ankyrin repeat protein YahD | | 0.42 | -2.81 |
| P02920 | Lactose permease (Lactose-proton symport) | | 0.53 | -2.36 |
| Q47539 | Taurine transport system permease protein TauC | Sulfur metabolism, ABC transporters | 0.48 | -2.81 |
| P0AAR5 | Inner membrane protein YbaN | | 0.42 | -2.81 |
| P0AFW4 | Regulator of nucleoside diphosphate kinase | | 0.40 | -2.61 |
| P0AER5 | Glutamate/aspartate import permease protein GltK | ABC transporters, Two-component system | 0.50 | -2.81 |
| P0AC44 | Succinate dehydrogenase hydrophobic membrane anchor subunit | Citrate cycle (TCA cycle), Butanoate metabolism, Oxidative phosphorylation | 0.48 | -5.14 |
| P0AAX6 | Uncharacterized protein McbA (MqsR-controlled colanic acid and biofilm protein A) | | 0.41 | -2.81 |
| P0AA67 | Threonine/homoserine exporter RhtA | | 0.42 | -2.52 |
| P0AAZ0 | Inner membrane protein YbjO | | 0.38 | -2.81 |
| P75917 | Uncharacterized protein YmdA | | 0.39 | -2.18 |
| P0ABX2 | Flagellar basal-body rod protein FlgC (Putative proximal rod protein) | Flagellar assembly | 0.38 | -3.81 |
| P75968 | Uncharacterized protein YmfE | | 0.35 | -2.81 |
| P11350 | Respiratory nitrate reductase 1 gamma chain (EC 1.7.5.1) (Cytochrome B-NR) (Nitrate reductase A subunit gamma) (Quinol-nitrate oxidoreductase subunit gamma) | Nitrogen metabolism, Two-component system | 0.41 | -3.16 |
| P0AFH6 | Oligopeptide transport system permease protein OppC | Quorum sensing, ABC transporters | 0.37 | -3.61 |
| P0ACW4 | Uncharacterized protein YdcA | | 0.43 | -2.81 |
| P76169 | UPF0060 membrane protein YnfA | | 0.43 | -2.81 |
| P76180 | Inner membrane protein YdgK | | 0.47 | -2.81 |
| P77389 | Inner membrane transport protein YdhP | | 0.47 | -3.31 |
| P76236 | Probable diguanylate cyclase CdgI (DGC) (EC 2.7.7.65) | | 0.48 | -3.07 |
| P69805 | PTS system mannose-specific EIID component (EII-M-Man) (EIID-Man) (Mannose permease IID component) | Phosphotransferase system (PTS), Fructose and mannose metabolism, Amino sugar and nucleotide sugar metabolism | 0.43 | -2.34 |
| P76264 | Probable manganese efflux pump MntP | | 0.51 | -2.81 |
| P33219 | Protein YebF | | 0.37 | -2.39 |
| P36561 | Adenosylcobinamide-GDP ribazoletransferase (EC 2.7.8.26) (Cobalamin synthase) (Cobalamin-5'-phosphate synthase) | Porphyrin metabolism | 0.36 | -2.41 |
| P33015 | UPF0394 inner membrane protein YeeE | | 0.41 | -2.58 |
| P64536 | Uncharacterized protein YeiS | | 0.41 | -2.81 |
| P33021 | Putative nucleoside permease NupX | | 0.39 | -2.81 |
| P0ABW3 | Uncharacterized ferredoxin-like protein YfaE | | 0.42 | -2.81 |
| P0AEB0 | Sulfate transport system permease protein CysW | Sulfur metabolism, ABC transporters | 0.49 | -2.49 |
| P45956 | CRISPR-associated endoribonuclease Cas2 (EC 3.1.-.-) | | 0.45 | -2.81 |
| P0ADR2 | UPF0382 inner membrane protein YgdD | | 0.39 | -2.81 |
| P11667 | Arginine exporter protein ArgO | | 0.46 | -3.31 |
| P64606 | Intermembrane phospholipid transport system permease protein MlaE | ABC transporters | 0.42 | -3.31 |
| P45761 | Type II secretion system protein J (T2SS protein J) (Putative general secretion pathway protein J) | Bacterial secretion system | 0.44 | -2.81 |
| P09391 | Rhomboid protease GlpG (EC 3.4.21.105) (Intramembrane serine protease) | | 0.42 | -2.59 |
| P0A6V5 | Thiosulfate sulfurtransferase GlpE (EC 2.8.1.1) | Sulfur metabolism | 0.36 | -4.81 |
| P0AET5 | Protein HdeD | | 0.50 | -2.81 |
| P31456 | Protein CbrA (CreB-regulated gene A protein) | | 0.36 | -2.24 |
| P32139 | Uncharacterized protein YihR | Galactose metabolism, Glycolysis / Gluconeogenesis | 0.41 | -2.61 |
| P32166 | 1,4-dihydroxy-2-naphthoate octaprenyltransferase (DHNA-octaprenyltransferase) (EC 2.5.1.74) | Ubiquinone and other terpenoid-quinone biosynthesis ' | 0.41 | -2.69 |
| P32167 | Uncharacterized protein YiiX | | 0.38 | -2.33 |
| P0A7C8 | Protein PsiE | | 0.48 | -2.81 |
| P0A9E2 | Regulatory protein SoxS | | 0.42 | -2.63 |
| P0AFJ1 | Protein YjdM | | 0.36 | -2.59 |
| P39282 | Inner membrane transporter YjeM | | 0.50 | -2.81 |
| P0ADD9 | Uncharacterized protein YjjY | | 0.42 | -3.81 |
| P0ADB4 | Entericidin A | | 0.44 | -3.81 |
| P0AE26 | L-arabinose transport system permease protein AraH | ABC transporters | 0.44 | -3.81 |
| Q2EET2 | UPF0370 protein YpfN | | 0.44 | -2.81 |
| A5A628 | Uncharacterized protein YjbT | | 0.38 | -2.81 |
| P58036 | Uncharacterized protein YjbS | | 0.45 | -2.81 |

**Supplementary Table 7.** A list of proteins that overlap between M2D2 drug – protein interactions and CRISPRi negative log fold change genes. More specifically, this list is the overlap between hits by the drug fasudil (top 80$^{th}$ percentile of interaction scores) predicted by M2D2 and genes that, when knocked down by CRISPRi, resulted in loss of fitness. The M2D2 importance filter of 15% was not applied to the list of proteins unlike Supplementary Table 6. Full names of the proteins were taken from UniProt[5]. If available, the Kegg database[6] pathways that the protein is a part of are noted. The list is sorted by the largest total score between ML and the absolute value of the CRISPRi log fold change.

| Fosfomycin mechanism | | | | | | |
|---|---|---|---|---|---|---|
| **UniProt ID** | **gene** | **gene full name** | **Fasudil – protein interaction score** | **rank no ML filter (out of 4070)** | **Importance score** | **Importance score rank (out of 8140)** |
| P0AC78 | *wecA* | Undecaprenyl-phosphate alpha-N-acetylglucosaminyl 1-phosphate transferase (EC 2.7.8.33) (UDP-GlcNAc:undecaprenyl-phosphate GlcNAc-1-phosphate transferase) (Undecaprenyl-phosphate GlcNAc-1-phosphate transferase) | 0.413 | 478 | 0.028 | 718 |
| P0AGC0 | *uhpT* | Hexose-6-phosphate:phosphate antiporter | 0.402 | 527 | 0.007 | 1984 |
| P0A6W3 | *mraY* | Phospho-N-acetylmuramoyl-pentapeptide-transferase (EC 2.7.8.13) (UDP-MurNAc-pentapeptide phosphotransferase) | 0.454 | 290 | 0 | 5194 |

| ***cysB*** **mutation, peptidoglycan biosynthesis, and Tol-Pal complex mechanism** | | | | | | |
|---|---|---|---|---|---|---|
| **UniProt ID** | **gene** | **gene full name** | **Fasudil – protein interaction score** | **rank no ML filter (out of 4070)** | **Importance score** | **Importance score rank (out of 8140)** |
| P0ABH4 | *mreD* | Rod shape-determining protein MreD | 0.477 | 204 | 0.134 | 32 |
| P19934 | *tolA* | Tol-Pal system protein TolA | 0.3 | 1278 | 0.121 | 40 |
| P0ABV6 | *tolR* | Tol-Pal system protein TolR | 0.365 | 720 | 0.043 | 383 |
| P45955 | *cpoB* | Cell division coordinator CpoB | 0.439 | 353 | 0.029 | 699 |
| P27434 | *rodZ* | Cytoskeleton protein RodZ | 0.365 | 729 | 0.027 | 777 |
| P0AAN5 | *yaiA* | Uncharacterized protein YaiA | 0.356 | 782 | 0.027 | 780 |
| P0AFT2 | *tcyL* | L-cystine transport system permease protein TcyL | 0.437 | 360 | 0.019 | 1124 |

**Supplementary Table 8.** A list of proteins selected for high ML and importance scores, and their link to key mechanisms discussed in results section. There are 4070 *E. coli* proteins used in the M2D2 model. Each protein has both a sigma and delta score used as input into the second ML stage of M2D2 to predict drug – drug interactions. The fasudil – protein interaction scores infer the strength of interaction between drug and protein. The importance scores are based on the M2D2 drug combination predictions as a whole and are not fasudil specific. The table is sorted by importance score. The table is separated into two categories of mechanisms discussed in the main text. Gene names were taken from the UniProt[5] database.

| Stressor | Pearson’s r | p-value |
|---|---|---|
| PH4 | 0.74 | 1.08E-18 |
| NOREPINEPHRINE | 0.60 | 3.11E-11 |
| GLUFOSFOMYCIN | 0.60 | 4.52E-11 |
| SPECTINOMYCIN | 0.57 | 6.74E-10 |
| INDOLICIDIN | 0.57 | 7.01E-10 |
| FOSFOMYCIN | 0.54 | 6.60E-09 |
| MMS | 0.53 | 1.79E-08 |
| ETHANOL | 0.52 | 3.20E-08 |
| PH10 | 0.52 | 3.55E-08 |
| DOXORUBICIN | 0.51 | 4.31E-08 |
| GENTAMICIN | 0.51 | 6.44E-08 |
| NACL | 0.50 | 1.06E-07 |
| CCCP | 0.49 | 1.97E-07 |
| MITOMYCINC | 0.49 | 2.25E-07 |
| DOXYCYCLINE | 0.48 | 4.95E-07 |
| CEFSULODIN | 0.48 | 4.98E-07 |
| EGCG | 0.48 | 5.23E-07 |
| DIBUCAINE | 0.48 | 5.91E-07 |
| STREPTOMYCIN | 0.46 | 1.26E-06 |
| PH5 | 0.46 | 1.61E-06 |
| PHLEOMYCIN | 0.45 | 2.07E-06 |
| ACRIFLAVINE | 0.45 | 2.63E-06 |
| CECROPINB | 0.45 | 3.43E-06 |
| NORFLOXACIN | 0.43 | 8.74E-06 |
| PH6 | 0.42 | 1.26E-05 |
| NIGERICIN | 0.41 | 2.11E-05 |
| AMIKACIN | 0.41 | 2.76E-05 |
| AMOXICILLIN | 0.40 | 3.12E-05 |
| GLUCOSE | 0.40 | 3.37E-05 |
| BACITRACIN | 0.38 | 8.05E-05 |
| EGTA | 0.38 | 1.13E-04 |
| RADICICOL | 0.37 | 1.34E-04 |
| HIGHCOPPER | 0.37 | 1.46E-04 |
| ISONIAZID | 0.37 | 1.61E-04 |
| MINOCYCLINE | 0.37 | 1.78E-04 |
| CHOLATE | 0.36 | 1.89E-04 |
| SDS | 0.36 | 2.26E-04 |
| AZITHROMYCIN | 0.36 | 2.54E-04 |
| LEVOFLOXACIN | 0.35 | 3.20E-04 |
| BLEOMYCIN | 0.35 | 3.57E-04 |
| ANAEROBIC | 0.35 | 3.87E-04 |
| NALIDIXICACID | 0.34 | 5.00E-04 |
| PH8 | 0.34 | 5.39E-04 |

**Supplementary Table 9.** Top 45, out of 114, correlated stressors, from Nichols *et al.*[11] chemogenomic study, with fasudil M2D2 predictions from this study. P-values for Pearson’s r values were all less than 1E-8. The top stressors were chosen based on M2D2 correlations. 114 unique stressors, from the Nichols et al. dataset, were assessed.

| Stressor | Pearson's r | p-value |
|---|---|---|
| GLUFOSFOMYCIN | 0.66 | 1.0E-13 |
| PH4 | 0.63 | 1.6E-12 |
| SPECTINOMYCIN | 0.61 | 1.2E-11 |
| INDOLICIDIN | 0.61 | 1.8E-11 |
| EGCG | 0.59 | 7.1E-11 |
| FOSFOMYCIN | 0.57 | 4.6E-10 |
| ANAEROBIC | 0.55 | 3.3E-09 |
| CARBENICILLIN | 0.54 | 5.2E-09 |
| NOREPINEPHRINE | 0.54 | 5.9E-09 |
| DOXYCYCLINE | 0.54 | 8.0E-09 |
| PH6 | 0.54 | 8.4E-09 |
| DOXORUBICIN | 0.53 | 1.3E-08 |
| AZIDOTHYMIDINE | 0.53 | 1.4E-08 |
| CEFSULODIN | 0.51 | 4.8E-08 |
| PH10 | 0.50 | 1.2E-07 |
| CCCP | 0.50 | 1.5E-07 |
| GLUCOSE | 0.48 | 3.3E-07 |
| CEFOXITIN | 0.48 | 3.5E-07 |
| SULFAMONOMETHOXINE | 0.47 | 6.6E-07 |
| PH8 | 0.47 | 6.9E-07 |
| CHOLATE | 0.47 | 7.1E-07 |
| MMS | 0.47 | 9.2E-07 |
| NIGERICIN | 0.46 | 1.1E-06 |
| PROCAINE | 0.46 | 1.5E-06 |
| NACL | 0.46 | 1.7E-06 |
| PHLEOMYCIN | 0.46 | 1.8E-06 |
| AZTREONAM | 0.45 | 2.4E-06 |
| TAUROCHOLATE | 0.45 | 2.5E-06 |
| RADICICOL | 0.45 | 2.6E-06 |
| VERAPAMIL | 0.45 | 3.2E-06 |
| MECILLINAM | 0.44 | 3.9E-06 |
| DIBUCAINE | 0.44 | 4.1E-06 |
| SULFAMETHIZOLE | 0.44 | 4.9E-06 |
| RIFAMPICIN | 0.44 | 5.2E-06 |
| ERYTHROMYCIN | 0.44 | 5.2E-06 |
| AMIKACIN | 0.44 | 5.9E-06 |
| HYDROXYUREA | 0.43 | 6.4E-06 |
| ETHANOL | 0.43 | 7.5E-06 |
| EGTA | 0.43 | 7.9E-06 |
| ACTINOMYCIND | 0.43 | 8.5E-06 |
| PARAQUAT | 0.43 | 9.1E-06 |
| METHOTREXATE | 0.43 | 1.0E-05 |
| PH5 | 0.42 | 1.4E-05 |
| LEVOFLOXACIN | 0.42 | 1.5E-05 |
| A22 | 0.41 | 1.9E-05 |

**Supplementary Table 10.** Top 45, out of 114, correlated stressors, from Nichols *et al.*[11] chemogenomic study, with fasudil CRISPRi data from this study. P-values for Pearson's r values were all less than 1E-8. The top stressors were chosen based on M2D2 correlations. 114 unique stressors, from the Nichols et al. dataset, were assessed.

### SUPPLEMENTARY DISCUSSION

#### *cysB* mutation and Tol-Pal system

TcyL is a L-cystine transport system permease protein, associated with the ABC transporters pathway and also linked to mecillinam resistance[5–7]. YaiA is an uncharacterized protein, but it has also been shown to increase, like TcyL, due to the CysB resistance mutation[5,7]. Fasudil could be targeting proteins that would otherwise increase with mecillinam resistance mutations. Perhaps this decreases the bioavailability of proteins, which are involved in alternative metabolisms prompted by the *cysB* mutation. If fasudil is also attacking the sulfur metabolism of the cell, thereby decreasing the cysteine in the metabolic system, the potential cell response of increasing LpoB, PBP1B and associated proteins could still be thwarted by fasudil's targeting of TcyL and YaiA.

Recent literature also suggest the Tol-Pal system of gram-negative bacteria is closely linked to PBP1B-LpoB and the ability of the cell to mediate outer membrane constriction during cell division[8]. The Tol-Pal complex has also been linked to pathogenicity in many gram-negative bacteria, making it a promising target for antibiotic development[9]. CpoB, a key protein in the interactions between the Tol-Pal and PBP1B-LpoB, has a strong interaction with fasudil in our M2D2 model. The CpoB is the facilitator protein between peptidoglycan synthesis by PBP1B-LpoB and outer membrane constriction by the Tol-Pal system[9,10]. M2D2 also predicts fasudil to interact strongly with TolR and TolA, key proteins in the Tol-Pal complex. Thus, it appears that fasudil can disrupt the Tol-Pal system, which directly impacts the cell's peptidoglycan biosynthesis. Perhaps the Tol-Pal disruption also decreases PBP1B-LpoB functionality, thereby negating the effects of mecillinam resistance created by reduced cysteine production.

Overall, specific protein targets predicted by M2D2 seem to indicate that fasudil can facilitate mecillinam antimicrobial activity by attacking peptidoglycan biosynthesis both directly and indirectly through the Rod and Tol-Pal systems. Fasudil may also target specific proteins that are produced when cysteine biosynthesis is disrupted, possibly by sulfur metabolism disruption. This specific set of protein targets could prevent any conferred resistance to mecillinam due to decreased cysteine levels.

**REFERENCES**

1. Huang, K. *et al.* DeepPurpose: a deep learning library for drug–target interaction prediction. *Bioinformatics* 1–6 (2020).
2. Deng, Y. *et al.* A multimodal deep learning framework for predicting drug-drug interaction events. *Bioinformatics* **36**, 4316–4322 (2020).
3. Albo, J. *et al.* EZ-SPOTs: A simple and robust high-throughput liquid handling platform. *bioRxivorg* 2024.05.13.594031 (2024).
4. Shiri, S. *et al.* Surface Patterned Omniphobic Tiles (SPOTs): a versatile platform for scalable liquid handling. *bioRxiv* 2024.01.17.575712 (2024) doi:10.1101/2024.01.17.575712.
5. UniProt Consortium. UniProt: the universal protein knowledgebase in 2021. *Nucleic Acids Res.* **49**, D480–D489 (2021).
6. Kanehisa, M. & Goto, S. KEGG: kyoto encyclopedia of genes and genomes. *Nucleic Acids Res.* **28**, 27–30 (2000).
7. Thulin, E. & Andersson, D. I. Upregulation of PBP1B and LpoB in cysB mutants confers mecillinam (amdinocillin) resistance in Escherichia coli. *Antimicrob. Agents Chemother.* **63**, (2019).
8. Typas, A. *et al.* Regulation of peptidoglycan synthesis by outer-membrane proteins. *Cell* **143**, 1097–1109 (2010).
9. Hirakawa, H., Suzue, K. & Tomita, H. Roles of the Tol/pal system in bacterial pathogenesis and its application to antibacterial therapy. *Vaccines (Basel)* **10**, 422 (2022).
10. Gray, A. N. *et al.* Coordination of peptidoglycan synthesis and outer membrane constriction during Escherichia coli cell division. *Elife* **4**, (2015).
